# Cerebral Blood Volume Modulates Glymphatic Influx Through Extra-ventricular Cerebrospinal Fluid Volume

**DOI:** 10.1101/2024.01.15.575561

**Authors:** Juchen Li, Xingyue Liu, Binshi Bo, Mengchao Pei, Kaiwei Zhang, Chuanjun Tong, Ming Jiang, Sheng Zhang, Yufeng Li, Jing Cang, Zhifeng Liang, Fang Fang

## Abstract

The glymphatic system facilitates waste removal via cerebrospinal fluid (CSF) influx alongside perivascular spaces throughout the brain. Vasomotion, the slow motion of blood vessel (0.1-0.3 Hz), has been found to be one of the driving forces for perivascular clearance, but it is not clear whether more chronical change of vessel diameter, as reflected by macroscopic cerebral blood volume (CBV), has any impact on glymphatic function. Combining multimodal mouse MRI techniques, we investigated the relationship among glymphatic influx, CBV, CSF volume and EEG power under six different conditions (awake, dexmedetomidine, isoflurane, isoflurane/dexmedetomidine, ketamine/xylazine and awake with caffeine). We found dexmedetomidine and caffeine enhanced glymphatic influx, while isoflurane reduced it compared with awake condition. Quantitative CBV imaging revealed that glymphatic influx was negatively correlated to CBV across the above conditions. Furthermore, such negative correlation was found to be mediated in part by changes of extra-ventricular CSF volume, which was quantified using T1 MRI. Taken together, our results suggest that CBV is a consciousness independent modulator of glymphatic function and modulates glymphatic influx through extra-ventricular CSF volume. This new finding opens potential avenues to enhance brain waste clearance by regulating CBV, which could be beneficial for protein deposition related neurological diseases.

**Teaser:** CBV is a consciousness independent modulator of glymphatic function and modulates glymphatic influx through extra-ventricular CSF volume.

## Introduction

The glymphatic system, a recently discovered network, serves as a crucial mechanism for clearing brain waste, including amyloid-β (Aβ) and tau (*1, 2*). This intricate system involves the movement of cerebrospinal fluid (CSF) along perivascular space (PVS) and its exchange with interstitial fluid (*3, 4*). Previous studies have indicated a potential correlation between glymphatic function and consciousness levels. Specifically, it has been shown that CSF influx to the brain is higher in mice during sleep or anesthesia with ketamine/xylazine (KX) compared with awake condition (*5*). Conversely, isoflurane (ISO)-induced unconsciousness has been reported to inhibit glymphatic influx in comparison to the awake condition (*6*). Moreover, the dynamics of glymphatic flow appear closely linked to the physiological effects induced by anesthetics, such as alterations in heart rate and EEG delta wave power (*7*). Despite these observations, the exact relationship and underlying mechanisms linking glymphatic function to (un)consciousness remain elusive.

As the scaffolding of glymphatic system, the cerebral blood vessels play a pivotal role in propelling CSF flow. A previous mathematical modeling study postulated that cerebrovascular pulsation was the driving force that promoted perivascular tracer into the brain (*8*). Subsequent research on anesthetized mice supported this notion, demonstrating that dobutamine-induced arterial pulsation increased tracer flow into the brain (*9*). Additionally, a two-photon study revealed a correlation between the rate of fluorescence tracer decay and low-frequency vasomotion, with suppression observed when isoflurane (ISO) inhibited vasomotion (*10*). These findings collectively suggest a connection between glymphatic influx and the dynamics of pulsation and vasomotion. However, vascular diameters also change at much slower time scales (hours, days or even longer) due to physiological or pathological conditions, and impacts of such relatively long-term vasoconstriction or vasodilation on glymphatic influx remains unexplored.

Recent studies have provided insights into the interplay between cerebrovascular constriction and dilation and glymphatic influx. Anesthesia with isoflurane combined with dexmedetomidine (ISODEX) was found to enhance glymphatic transport and induce greater vasoconstriction in the straight sinus compared with ISO condition in an anesthetized rat MRI study (*11*). Additionally, cerebral arterial constriction induced by optogenetic stimulation and spreading ischemia increased the periarterial space, resulting in heightened CSF inflow (*12, 13*). Conversely, hypercapnia was observed to suppress the glymphatic influx, potentially due to its cerebral vasodilatory effect (*14*). These findings indicate the significance of cerebrovascular constriction and dilation in influencing glymphatic influx. However, the exact mechanisms of cerebrovascular constriction and dilation on glymphatic influx remain unclear. Considering that the cranial cavity is an enclosed space consisting of brain tissue, cerebral blood and CSF, alterations in cerebral blood volume (CBV) induced by cerebrovascular constriction and dilation could significantly impact CSF volume (*15*). Therefore, exploring whether changes in CBV can modulate glymphatic influx by influencing CSF volume is a promising direction.

Herein, to explore the relationship among CBV, CSF volume and glymphatic influx in mice, we utilized multi-modal MRI techniques including dynamic contrast-enhanced MRI (DCE-MRI), T1 mapping and quantitative CBV MRI in six conditions (awake, DEX, ISO, ISODEX, KX and awake with caffeine). We found that across six conditions, glymphatic influx was negatively associated with CBV through potential contribution of positively correlated extra-ventricular CSF volume. These results suggest that CBV serves as a consciousness-independent modulator of glymphatic influx. More importantly, the results established a link among CBV change, brain volume change, extra-ventricular CSF volume and glymphatic influx, presenting a novel perspective on glymphatic function modulation. The current findings may be of potential clinical relevance in neurological diseases, in which intervention on CBV may be beneficial through promoting glymphatic function.

## Results

### Anesthesia/sedation increased or decreased glymphatic influx respectively compared to awake condition

Taking advantage of our extensive experience in awake mouse MRI (*16-19*), we compared glymphatic function under awake, dexmedetomidine (DEX), isoflurane (ISO), isoflurane/dexmedetomidine (ISODEX), ketamine/xylazine (KX) conditions using DCE-MRI. Given known impact of physiological parameters on glymphatic function (*7, 14*), we optimized the DCE-MRI paradigm for awake and mechanically ventilated anesthetized mice, including chronic cistern magna (CM) cannulation, postoperative recovery, habituation training for awake imaging, and endotracheal intubation for anesthetized imaging (Fig. 1A and Fig. S1). Such optimized paradigm ensured that animals were under normal and stable conditions with regards to key physiological parameters (Fig. S2), including PO_2,_ PCO_2_, SaO_2_, rectal temperature, across all anesthesia states.

**Fig. 1.**
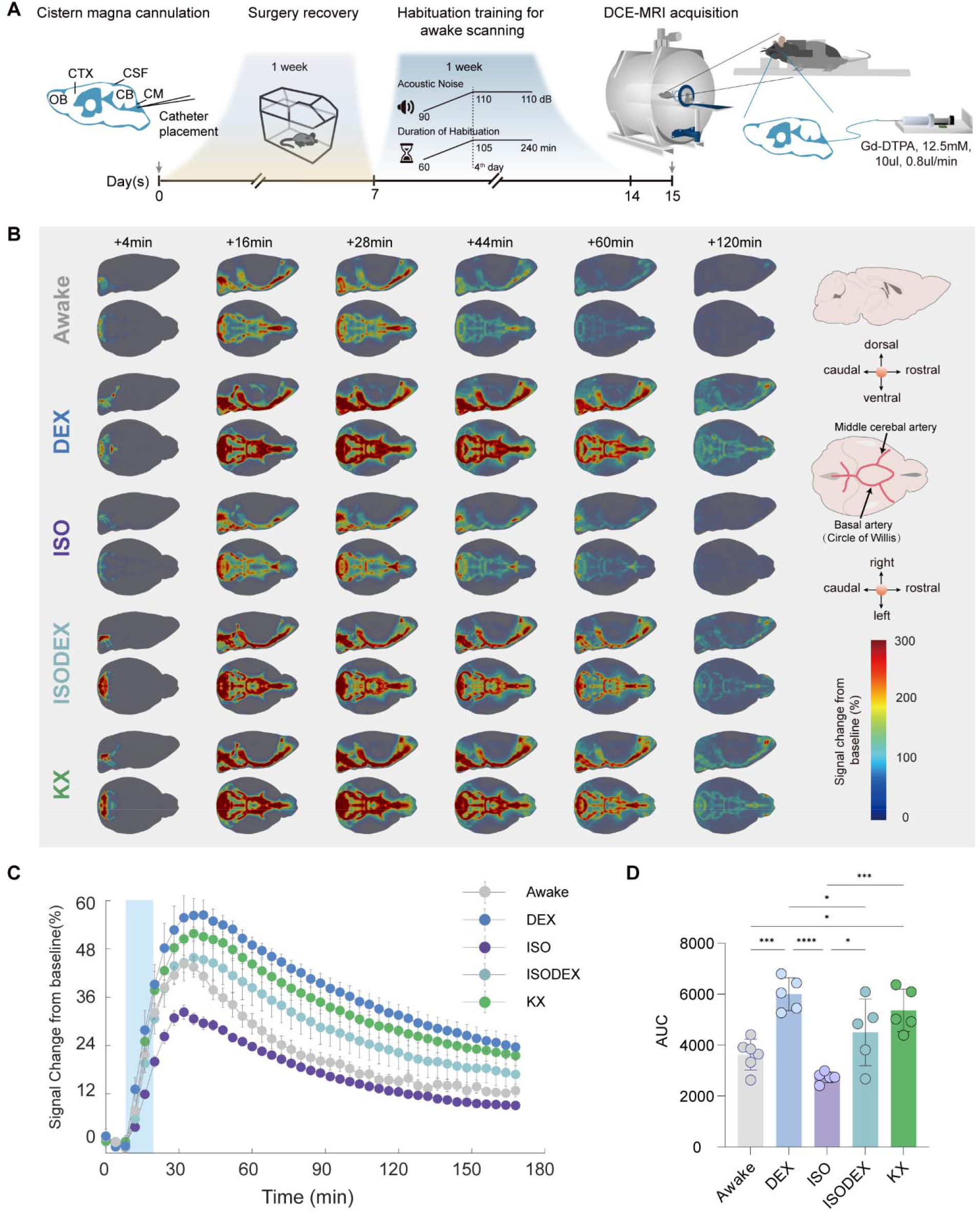
Gadolinium influx was modulated by anesthesia/sedation. **(A)** Schematic representation of the awake mouse DCE-MRI paradigm. **(B)** The three-dimensional (3D) spatiotemporal maps of averaged Gd-DTPA-induced signal changes from mice under awake (n = 6), dexmedetomidine (DEX, n = 5), isoflurane (ISO, n = 5), isoflurane/dexmedetomidine (ISODEX, n = 5) and ketamine/xylazine (KX, n = 5) conditions. Right panel, diagrams of the sagittal view (top) and ventral view (bottom) of the mouse brain. The black arrows pointed to the middle cerebral artery and basal artery (Circle of Willis). **(C)** The averaged time signal curves (TSCs) of Gd-DTPA-induced signal changes from whole brain of mice across the above five conditions. Data are presented as mean ± SEM. The blue rectangular area represented the contrast agent injection period (12.5 minutes). **(D)** Statistics of the area under curves (AUCs) of TSCs in five groups shown in **C**. One-way ANOVA with Tukey’s correction, *, p < 0.05; ***, p < 0.001; ****, p < 0.0001.

With the above paradigm, the spatiotemporal patterns of whole-brain glymphatic transport were obtained using gadolinium based DCE-MRI (Fig. 1B). Across all five conditions, gadolinium mainly flowed anteriorly along the Circle of Willis at the ventral brain surface, then proceeded dorsally along the middle cerebral artery, and moved toward the olfactory bulb, after intracisternally administering Gd-DTPA (Fig. 1B). Such pattern was highly consistent with previous reports (*7, 20*), which validated our current mouse DCE-MRI paradigm.

Among the five conditions, we observed increased DCE-MRI signals under DEX, ISODEX and KX conditions, while decreased signals under ISO were observed, all relative to the awake condition (Fig. 1, B, C and D). Such difference started to emerge early after Gd injection, and became prominent at ∼28 min after Gd injection. Quantitative analysis of the total brain DCE-MRI signals, as measured by area under curve (AUC, representing glymphatic transport over a 160min period), demonstrated that compared to awake mice (AUC: 3631 ± 603), mice under DEX (AUC: 6003 ± 646) and KX (5363 ± 838) exhibited a 65.32% and 47.70% increase respectively, while mice under ISO (AUC: 2743 ± 213) showed a 24.46% decrease (Fig. 1D). The whole-brain spatiotemporal patterns of DCE-MRI were illustrated in color-coded maps at five different time points (16 min, 28 min, 44 min, 60 min and 120 min) after Gd infusion (Fig. S3), and further quantified by calculating the number of voxels with significantly enhanced signal over time across five conditions (Fig. S4). These results clearly demonstrated that ISO reduced glymphatic influx while DEX and KX conditions increased the influx, compared to the awake condition.

### Cerebral blood volume was negatively correlated to glymphatic influx across awake and four anesthesia/sedation conditions

Based on the above DCE-MRI based whole-brain glymphatic transport patterns, we noted prominent signal changes along basal artery (Fig. 1B). DCE-MRI signals of basal artery (Fig. 2 A) showed a similar trend across five conditions (Fig. 2, B and C) as to the whole-brain signals (Fig. 1, C and D). At 28 minutes after injecting Gd-DTPA when the difference was prominent, DCE-MRI maps revealed that, compared to the awake condition, the magnitude and extent of signal enhancement in the regions surrounding the basal artery were higher under DEX and KX, and lower under ISO (Fig. 2, D to H). We further evaluated the relationship between the distance from basal artery (Fig. 2I) and the signal enhancement, and as expected, both signal amplitude and signal ascending slope showed strong dependence on the distance from basal artery across all five conditions (Fig. 2, J and K).

**Fig. 2.**
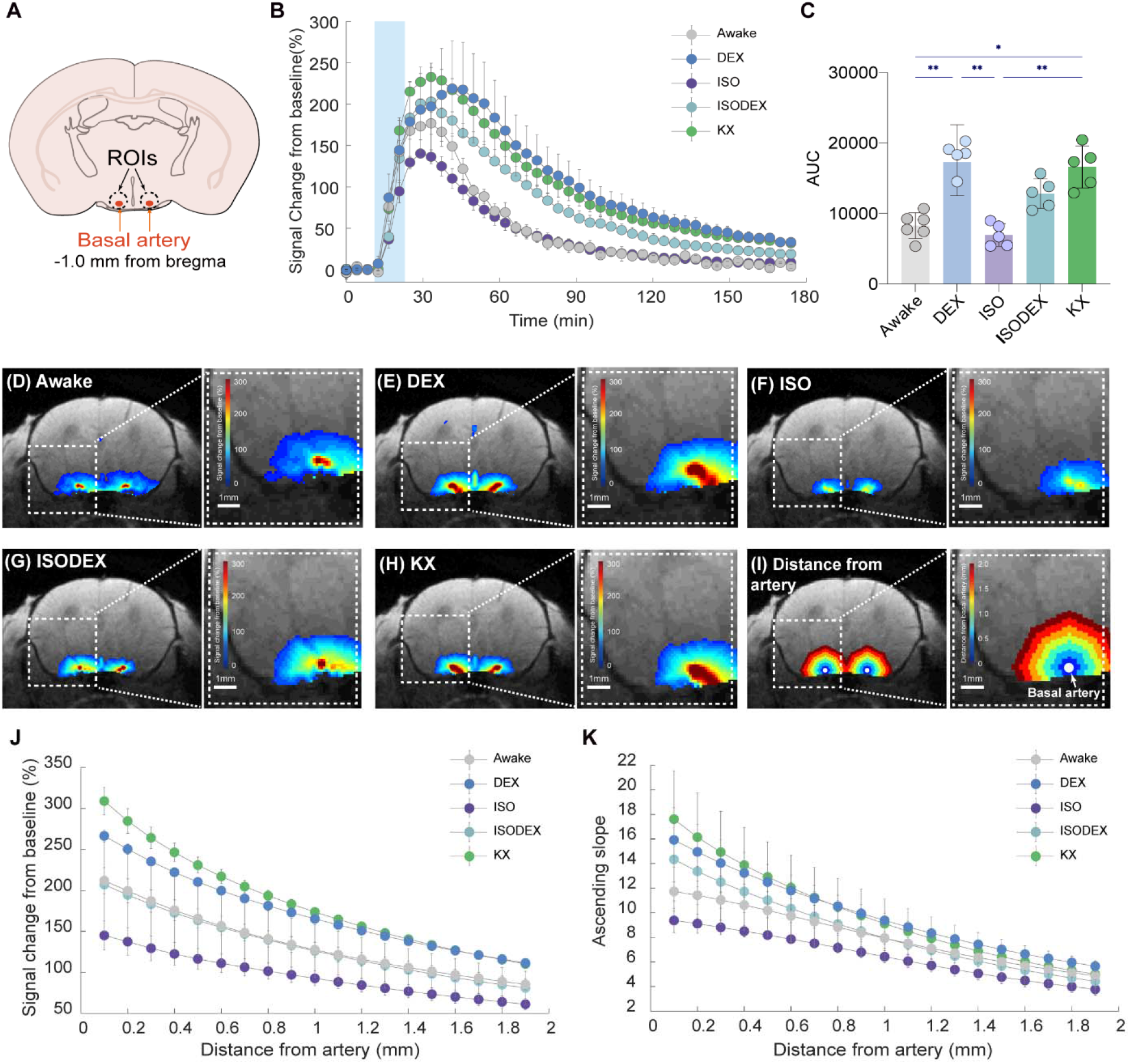
Glymphatic influx affected by anesthesia/sedation was prominent in the regions surrounding basal artery. **(A)** Schematic diagram of a coronal section (−1.0 mm from bregma of mouse brain) showing part of the basal artery. Dotted circles, the regions of interest (ROIs) surrounding the basal artery. **(B)** The TSCs of Gd-DTPA-induced signal changes from ROIs shown in **(A)** across five conditions. Data are presented as mean ± SEM. The blue rectangular area represented the contrast agent injection period (12.5 minutes). **(C)** Statistics of the AUCs of TSCs in five groups shown in **B**. One-way ANOVA with Tukey’s correction, *, p < 0.05, **, p < 0.01. **(D-H)** Averaged pseudocolour-coded maps (coronal section, -1.0 mm from bregma) of Gd-DTPA-induced signal changes 28 min after injecting Gd-DTPA in the awake **(D)**, DEX **(E)**, ISO **(F)**, ISODEX **(G)** and KX **(H)** conditions. Dotted boxes represented magnified images. **(I)** Distance from basal artery ranging from 0 to 2.0 mm in the coronal section (−1.0 mm from bregma). Dotted boxes represented magnified images. The white arrow pointed to the basal artery. **(J)** The averaged signal changes of different distances from basal artery 28 min after injecting Gd-DTPA across five conditions. **(K)** The averaged ascending slopes of signal change of different distances from basal artery after injecting Gd-DTPA across five conditions. Data are presented as mean ± SEM.

The above results, combined with the known vascular effect of DEX (vasoconstriction) and ISO (vasodilation), lead us to hypothesize that cerebral vasoconstriction (or vasodilation) increases (or decreases) CSF influx into the brain parenchyma. To assess the hypothesis, we utilized quantitative cerebral blood volume (CBV) imaging method, the “Bookend” MRI technique (*21-23*) (Fig. S5), to quantify CBV across awake and four anesthesia/sedation conditions (DEX, ISO, ISODEX and KX). Compared with awake group, the whole-brain CBV was decreased in the DEX, ISODEX and KX group and increased in the ISO group, especially in regions around large cerebral vessels such as basal arteries and superior sagittal sinus (Fig. 3A and Fig. S6A).

**Fig. 3.**
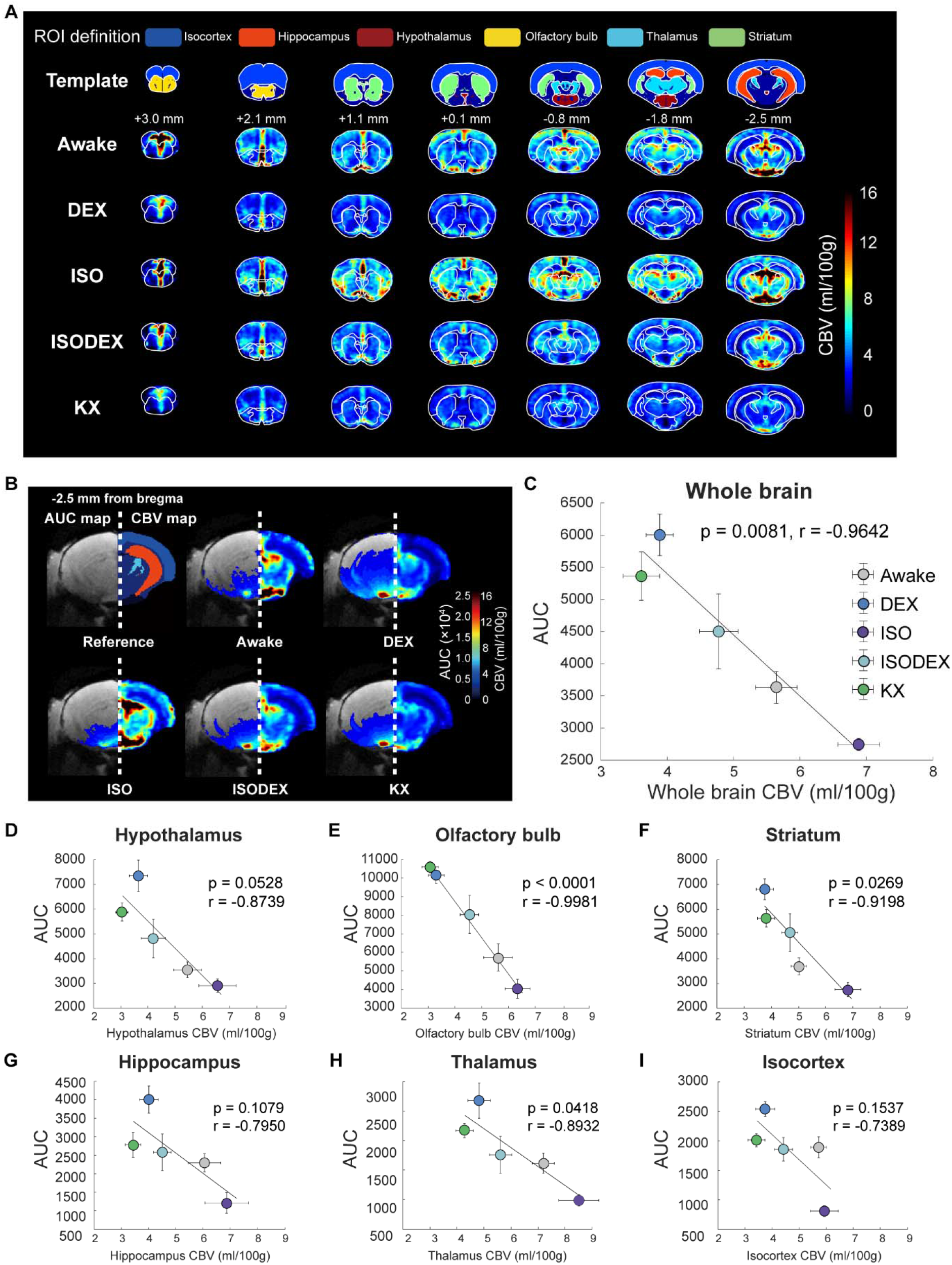
Cerebral blood volume was negatively correlated to glymphatic influx. **(A)** Whole brain CBV maps in awake (n = 9), DEX (n = 8), ISO (n = 6), ISODEX (n = 7) and KX (n = 6) conditions. Top row, coronal sections of the template showing six brain regions. Numbers under each slice denoted the distance from bregma. **(B)** Population-averaged pseudocolor-coded coronal images (–2.5 mm from bregma) of the Gd-DTPA induced AUC maps (AUC: cumulative value of signal change from 0 to 160min) (left, data from DCE-MRI images) and CBV maps (right). **(C)** Correlation analysis between averaged AUCs of TSCs and averaged CBV values from whole brain of five groups. **(D-I)** Correlation analysis between averaged AUCs of TSCs and averaged CBV values from hypothalamus (**D**), olfactory bulb **(E)**, striatum **(F)**, hippocampus **(G)** and thalamus **(H)** and isocortex **(I)** of five groups. Each dot represented the group average (whiskers, SEM).

Around the basal arteries, upon visual inspection we observed an association between greater overall tissue penetration of Gd-DTPA and lower CBV across the five conditions (Fig. 3B). To further determine the relationship between CBV and glymphatic influx, we conducted correlation analysis between the AUCs and CBV values across the whole brain, thalamus, striatum, olfactory bulb, hippocampus, hypothalamus, and isocortex in the five groups (Fig. S7). Correlation analysis revealed that glymphatic influx had a significant negative relationship with CBV in whole brain, thalamus, olfactory bulb, and striatum (Fig. 3, C to I). In addition, we found that peak amplitude of signal change and CBV were also negatively correlated (Fig. S6, B to I). Therefore, our findings suggested that as CBV increased (or decreased), glymphatic influx decreased (or increased) correspondingly, providing support for our hypothesis.

### Extra-ventricular CSF volume, which changed negatively with CBV, was positively correlated to glymphatic influx across awake and four anesthesia/sedation conditions

Next, we investigated that how the changes in CBV were linked to glymphatic influx. We hypothesized that changes in CBV could alter brain tissue volume, further influencing CSF volume given the fixed total cranial volume. Therefore, we further assessed CSF and brain tissue volume for exploring the relationship with CBV and glymphatic influx. Utilizing quantitative T1 MRI mapping method as CSF T1 is longer than brain tissue, we obtained whole-brain CSF probability maps under above five conditions (Fig. 4A and Fig. S8). These maps were then further segmented into ventricular and extra-ventricular components (Fig. 4B and Fig. S9). Compared with awake group, whole-brain CSF probability increased in DEX and KX groups while significantly decreased in ISO group, which was consistent with previous studies (*24*) (Fig. 4C and Fig. S10). Specifically, the regions showing prominent changes in the CSF probability maps were primarily in the CSF cisterns near cerebral blood vessels and the subarachnoid space (Fig. 4C and Fig. S10). Based on the CSF probability maps, we further quantified the CSF volume and found that changes in whole-brain CSF volume across the five conditions were consistent with the CSF probability maps (Fig. 4D). Interestingly, this change was mainly attributed to a significant difference in extra-ventricular CSF volume (Fig. 4E), as no significant difference in ventricular CSF volume among groups was observed (Fig. 4F).

**Fig. 4.**
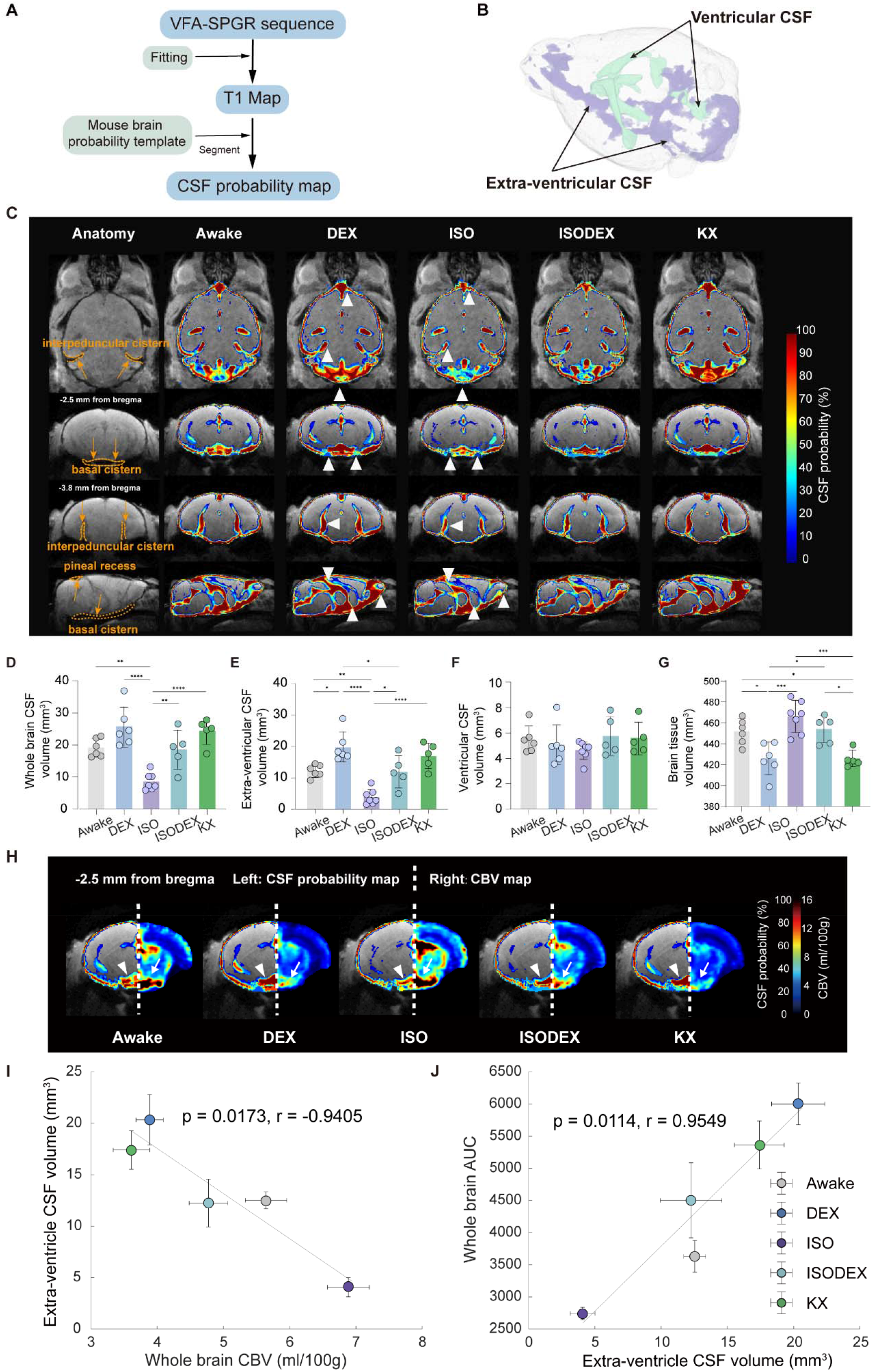
Glymphatic transport was positively correlated to the extra-ventricular CSF volume. **(A)** Flowchart for probabilistic segmentation of CSF volume. **(B)** 3D-reconstruction of T1 map showed two main CSF compartments: intra-ventricular CSF and extra-ventricular CSF. **(C)** Averaged whole-brain CSF probability maps from three views in awake (n = 6), DEX (n = 6), ISO (n = 7), ISODEX (n = 5) and KX (n = 5) group. Arrowheads denoted the regions with large changes in the CSF probability maps. The locations of CSF cisterns were marked with yellow arrows and dashed lines on the template images. **(D-G)** Statistical comparisons of whole brain CSF volume **(D)**, extra-ventricular CSF volume **(E)**, ventricular CSF volume **(F)** and brain tissue volume **(G)** in five groups. One-way ANOVA with Tukey’s correction, *, p < 0.05; **, p < 0.01; ***, p < 0.001; ****, p < 0.0001. **(H)** Pseudocolor-coded coronal images (–2.5 mm from bregma) of the averaged CSF probability maps (left) and CBV maps (right), designated by white arrows (CBV map) and arrowheads (CSF probability map) respectively. **(I)** Correlation analysis between whole brain CBV and extra-ventricular CSF volume across five groups. **(J)** Correlation analysis between extra-ventricular CSF volume and AUCs of whole brain TSCs across five groups. Each dot represented the group average (whiskers, SEM).

Based on T1 map, we also observed that changes in brain tissue volume were consistent with the changes in CBV (Fig. 4G), with higher CBV corresponding to larger brain volume (Fig. S11A), and an inverse relationship between brain tissue volume and whole CSF volume were also observed (Fig. S11B). We further observed an association between higher CSF probability and a lower CBV, particularly around the CSF cisterns, which were the part of the extra-ventricular CSF component (Fig. 4H). To further examine the relationships among glymphatic influx, CBV, and extra-ventricular CSF volume, we conducted correlation analyses among these three components. A significant negative correlation was observed between extra-ventricular CSF volume and whole-brain CBV (Fig. 4I), while extra-ventricular CSF volume showed a positive correlation with glymphatic influx (Fig. 4J).

To control for the potential impact of CSF probability threshold, two different CSF probability thresholds (0.8 and 0.9) were further applied to calculate CSF volume and performed correlation analyses (Fig. S12). Across two CSF probability thresholds, we consistently found a negative correlation between extra-ventricular CSF volume and CBV, and a positive correlation between extra-ventricular CSF volume and glymphatic influx. These results further supported the notion that extra-ventricular CSF volume could serve as a key link in the interaction between CBV and glymphatic influx.

### Caffeine reduced CBV and enhanced glymphatic influx

Next, we went on to examine the effect of caffeine on glymphatic function, as it is a known vasoconstrictor but does not induce anesthesia or sedation (*25*). As expected, EEG recordings demonstrated similar spectrograms and power spectrum density between the caffeine and awake groups, while DEX-sedated mice exhibited higher delta power (Fig. 5A). Furthermore, we observed that, compared with awake group, caffeine could reduce whole-brain CBV and brain tissue volume, while concurrently increasing extra-ventricular CSF volume, similar to the effects seen with DEX (Fig. 5, B to E).

**Fig. 5.**
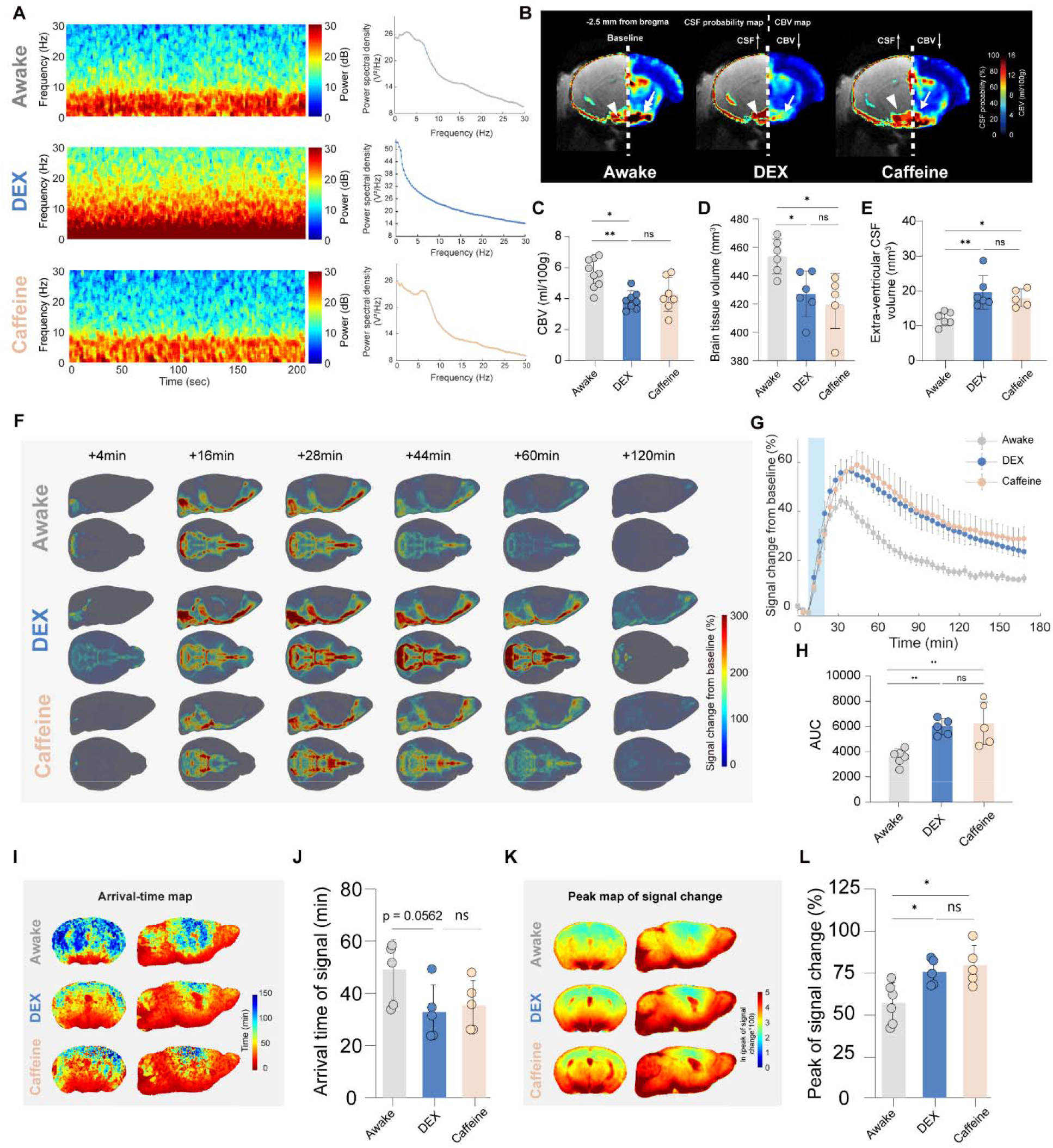
Caffeine reduced CBV and enhanced glymphatic influx. **(A)** Representative spectrogram of electroencephalogram (EEG, left panel) and corresponding power spectral density (PSD, right panel) from awake, DEX and caffeine groups, respectively. **(B)** Pseudocolor-coded coronal images (–2.5 mm from bregma) of the averaged CSF probability maps (left) and CBV maps (right) from awake, DEX and caffeine groups. Arrows and arrowheads denoted the regions with large changes in the CBV and CSF probability maps. **(C)** Statistical comparisons of whole brain CBV under three conditions (awake: n = 9, DEX: n = 8, caffeine: n = 7). Both caffeine and DEX significantly decreased whole brain CBV compared to awake group. **(D-E)** Statistical comparisons of brain tissue volume **(D)** and extra-ventricular CSF volume **(E)** under three conditions (awake: n = 6, DEX: n = 6, caffeine: n = 5). Both caffeine and DEX decreased brain tissue volume and increased extra-ventricular CSF volume compared to awake group. **(F)** The 3D spatiotemporal maps of averaged Gd-DTPA-induced signal changes in the awake (n = 6), DEX (n = 5), and caffeine (n = 5) groups. **(G)** The averaged TSCs of Gd-DTPA induced signal changes from whole brain of awake, DEX and caffeine groups. Data are presented as mean ± SEM. The blue rectangular area represents the contrast agent injection period (12.5 minutes). **(H)** Statistics of AUCs of TSCs in three groups shown in **G. (I)** Averaged arrival time maps of DCE-MRI signals in the awake, DEX and caffeine groups. **(J)** Statistics of arrival time from whole brain of awake, DEX and caffeine groups. **(K)** Peak maps of signal change in the awake, DEX and caffeine groups. **(L)** Statistics of peak values from whole brain of awake, DEX and caffeine groups. One-way ANOVA with Tukey’s correction, *, p < 0.05; **, p < 0.01; ns, no significance.

Importantly, caffeine also enhanced glymphatic influx. Both caffeine and DEX showed higher amplitude of signal change compared to the awake condition (Fig. 5, F to H). The 3D color-coded arrival time maps revealed that the Gd-DTPA took a relatively short time to reach the whole brain in the caffeine and DEX groups (Fig. 5, I and J), suggesting less resistance for the contrast agent influx into brain tissue. Additionally, the peak maps of signal change indicated that more Gd-DTPA entered brain tissue and covered a larger area in the caffeine and DEX groups (Fig. 5K). Compared with awake condition, peak values of signal change in whole brain were higher in caffeine and DEX group (Fig. 5L), suggesting that caffeine could enhance glymphatic function. Regional statistics were shown in Fig. S13, which were consistent with the whole brain results.

Considering that EEG activity was reported to be closely related to glymphatic function (*7*), we also analyzed the relationship between glymphatic influx and each EEG band wave, combining awake, caffeine, and four anesthesia/sedation conditions. However, no significant correlation was found between the peak values of whole-brain signal change and the ratios of delta, alpha, beta, and theta wave power to full band (1-20 Hz) power across all six conditions (Fig. S14). This result further implied that the regulation of glymphatic influx was independent of the consciousness level.

### CBV modulated glymphatic influx by influencing extra-ventricular CSF volume

Furthermore, we also conducted correlation analyses among glymphatic influx, CBV and extra-ventricular CSF volume across awake, caffeine, and four anesthesia/sedation conditions. Correlation analysis revealed that the negative correlation between CBV and glymphatic influx persist across above six conditions (Fig. 6A). In addition, a significant negative correlation was observed between extra-ventricular CSF volume and whole-brain CBV, while extra-ventricular CSF volume showed a positive correlation with glymphatic influx under six conditions (Fig. 6, B and C). The above results indicated an essential role of extra-ventricular CSF volume in mediating the relationship between CBV and glymphatic influx. To explore this, we further conducted a mediation analysis among CBV, glymphatic influx, and extra-ventricular CSF volume (Fig. 6D). We first demonstrated that the reduced CBV positively contributed to the increased AUC, which represents the glymphatic influx (total effect: regression coefficient c = - 870.067 ± 240.547, p < 0.001; Fig. 6D). Then, we incorporated the extra-ventricular CSF volume as a mediator. The direct relationship between the reduced CBV and the increased glymphatic influx became insignificant (direct effect: regression coefficient c’ = −257.313 ± 258.812, no significance) which meant a full mediation effect of the increased extra-ventricular CSF volume to the enhanced glymphatic influx. The ratio of the mediating effect to the total effect was 70.426%. This analysis indicated that CBV contributed negatively to glymphatic influx by exerting an influence on extra-ventricular CSF volume.

**Fig. 6.**
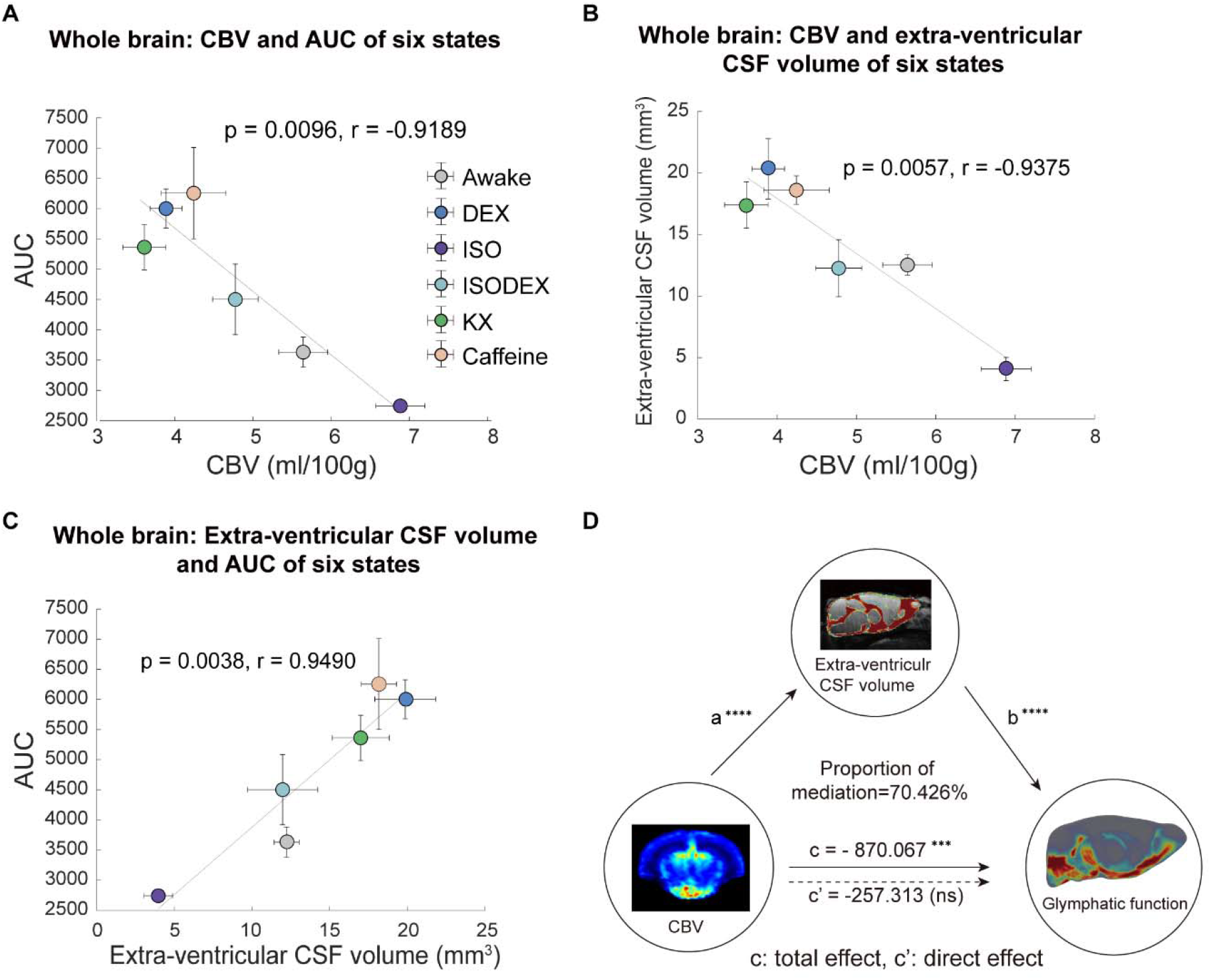
CBV negatively influences glymphatic influx through changes of extra-ventricular CSF volume. **(A)** Correlation analysis between averaged CBV values and averaged AUCs of TSCs from whole brain of six groups. **(B)** Correlation analysis between averaged CBV values and averaged extra-ventricular CSF volume from whole brain of six groups. **(C)** Correlation analysis between averaged extra-ventricular CSF volume and averaged AUCs of TSCs from whole brain of six groups. Each dot represented the group average (whiskers, SEM). **(D)** Mediation analysis among whole-brain CBV, glymphatic influx and extra-ventricular CSF volume. Mediation analysis was performed to evaluate the effect of CBV on extra-ventricular CSF volume (regression coefficient, a), the effect of extra-ventricular CSF volume on glymphatic function (regression coefficient, b), total effect (regression coefficient, c) of global CBV on glymphatic function and the direct effect (regression coefficient, c’) of global CBV on glymphatic function when extra-ventricular CSF volume was incorporated as a mediator. ***, p < 0.001; ****, p < 0.0001, ns, no significance.

Based on the above results, we revealed a CBV based mechanism of glymphatic modulation (Fig. 7), in which cerebral blood volume changes (vasoconstriction or vasodilation) regulate extra-ventricular CSF volume by reducing or increasing brain tissue volume, further influencing glymphatic influx.

**Fig. 7.**
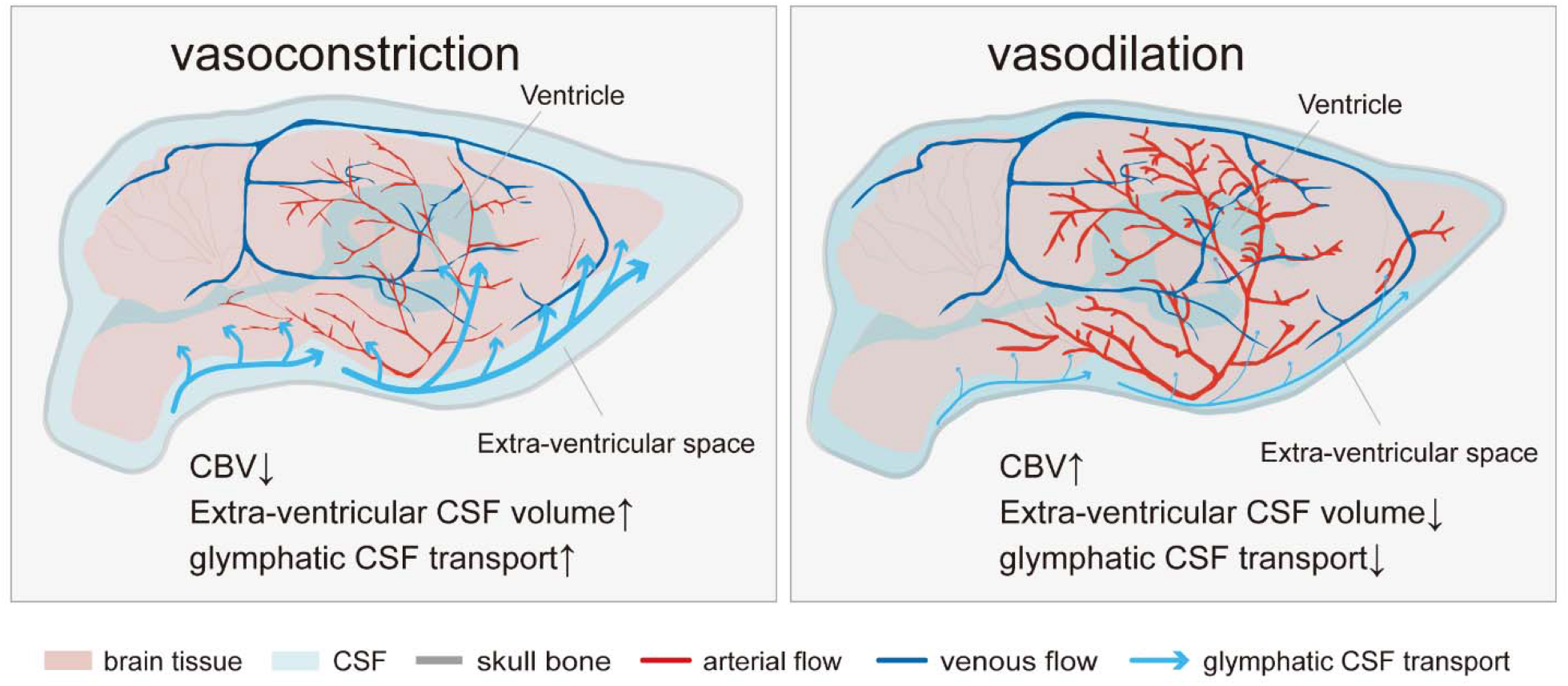
Schematic illustration of cerebral blood volume modulation of glymphatic influx. Vasoconstriction decreases whole-brain CBV, which increases the extra-ventricular CSF volume by altering the brain tissue volume, and enhances CSF influx into the brain parenchyma. Vasodilation increases CBV and decreases extra-ventricular CSF volume, which leads to less CSF influx into the brain parenchyma.

## Discussion

In the current study we explored the relationship between CBV and glymphatic function in mice based on a series of MRI techniques under six conditions, including awake, DEX, ISO, ISODEX, KX and awake with caffeine. Our results showed that glymphatic influx was negatively correlated to CBV, and such effect was mediated in part by changes of extra-ventricular CSF volume. Therefore, the current study indicates that CBV is a consciousness-level independent modulator of glymphatic function, and reveals a new CBV based mechanism of glymphatic modulation.

The glymphatic system plays a crucial role in clearing metabolic wastes from brain, and how it is regulated is of great interest and importance. Previous studies have examined the contribution of arterial pulsation (*9*), venous pulsation (*26, 27*) and slow vasomotion (*10*), sleep-wake cycle (*5, 28*) and certain anesthetic regimes (*7*) on glymphatic function. Interestingly, CBV and arterial diameters are also changed in above conditions (*29-32*), which suggests that cerebrovascular condition is closely related to glymphatic function.

CBV is well suited to potentially influence glymphatic influx. The cranial cavity is a fixed space, consisting of three components (cerebral blood, CSF and brain tissue). Therefore, volume changes of one component will need to be compensated by volume changes of other components or by inflow/outflow of CSF to spinal cord. Our current results demonstrated that across six conditions, CBV was negatively correlated to CSF volume, especially extra-ventricular CSF volume (Fig. 6B), which is in accordance with the Monroe-Kellie doctrine (*33*). A mouse 3DISCO study showed that in awake mice the CSF tracer flowed to a greater extent towards the spinal cord, while the CSF tracer was concentrated in the brain in KX anesthetized mice (*34*).

Their study provides additional support to the current result, as the reduced CBV and brain tissue volume could drive CSF from the spinal cord to the brain to maintain the normal intracranial pressure (ICP). Here, we did observe that extra-ventricular CSF volume was subjected to CBV change and positively related to the glymphatic function (Fig. 6, B and D), while the ventricular CSF volume remained relatively stable.

Importantly, mediation analysis further suggested that such extra-ventricular CSF mediated the CBV modulation effect of glymphatic function (Fig. 6D). CSF flows from the ventricular system into the subarachnoid space (extra-ventricular space), and mainly exits along basal cistern via the cranial/spinal nerves and arachnoid granulations (*35*). It is reasonable that increased extra-ventricular volume could decrease the resistance of CSF flow, which accelerates the CSF circulation. As a part of CSF circulation, glymphatic transport could be enhanced by the decreased resistance in CSF circulation, which was supported by the shorter arrival time under DEX and caffeine conditions compared with awake condition (Fig. 5, I and J). Previous studies have also reported that ISODEX increased CSF volume and enhanced glymphatic influx compared with ISO condition (*11, 24*), which is consistent with our results.

Other physiological conditions, for example sleep (*5*), have been reported to modulate glymphatic function. Interestingly, sleep has also been reported to exhibit various volumetric alterations, such as extracellular space (*5*) and CSF (*36*). A human study has observed that the CSF volume was increased during sleep based on voxel-based morphometry (VBM) analysis. Therefore, increased CSF volume during sleep may in part contribute to enhance glymphatic function, along with other factors, such as extracellular space fraction.

It is interesting to note that the current results, combining with the above sleep study, suggest that the regulation of the glymphatic system by CBV may be independent of consciousness level. It was reported that the enhancement of glymphatic function during sleep or anesthetized conditions was associated with higher slow delta wave power, which was typically observed during NREM sleep and certain anesthetic regiments (*5, 7, 11*). Therefore, it has been speculated that glymphatic function depends on consciousness level. The unique strength of the current study is that we included 6 different conditions, including 4 anesthesia/sedation (DEX, ISO, ISODEX and KX groups) and 2 awake conditions (awake and caffeine groups). Using the awake group as a baseline, our results clearly showed anesthesia/sedation and caffeine could either increase or decrease glymphatic influx. It is important to note that caffeine and DEX groups, both with similar vasoconstrictive effect but under the opposite consciousness conditions (sedated v.s. alert), exhibited similar increase of glymphatic influx (Fig. 4). Taken together, our results indicated that consciousness level per se is unlikely to be a direct modulator of glymphatic function, but rather other physiological factors (e.g., CBV) associated with consciousness level could be more direct modulators.

MRI is a set of powerful imaging tools, and it can be used to examine various aspects of the glymphatic system in addition to widely used DCE-MRI. In the current study, we employed a quantitative CBV mapping method (*21-23*) to quantify the quantitative CBV across various conditions. Furthermore, T1 mapping method was utilized for quantification of CSF volumes. Noninvasive whole brain mapping of CBV and CSF volume provided key measurements of current glymphatic modulation mechanism (Fig. 6), which would be difficult to achieve using other imaging modalities. Furthermore, ventricular CSF flow velocity can also be quantified by a novel phase-contrast MRI technique developed by us (*18*). Therefore, we believe that multimodal MRI techniques have great potential for further studying the functions and mechanisms of glymphatic system.

Despite its wide application in glymphatic studies, almost all animal DCE-MRI experiments were conducted under free-breathing anesthetized conditions with potentially abnormal physiological changes, such as hypercapnia. Importantly, hypercapnia has been reported to inhibit glymphatic influx (*14*). Therefore, it is critical to consider the physiological condition when examining glymphatic function using DCE-MRI. In addition, DCE-MRI studies in awake animals are sparse, which would serve as an essential baseline when comparing different conditions. Thus, in the current study we took advantage of our extensive awake mouse fMRI experience (*16-19*) and modified the setup to allow awake DCE-MRI. Furthermore, for anesthetized imaging, we intubated and mechanically ventilated the mice, with ventilator parameters carefully adjusted to achieve normal blood gas (Fig. S2). Therefore, the current study presented a set of highly optimized awake and anesthetized mouse MRI setups for glymphatic study.

There are several limitations in the current study. First, although the above findings have been obtained based on our multimodal MRI techniques, MRI cannot achieve the spatial resolution at the cellular level. Compared to MRI, optical methods such as two-photon imaging can provide higher spatiotemporal resolution for investigating glymphatic system, but with a much-limited spatial coverage and often only in superficial part of the cortex. Therefore, two types of imaging methods (MRI and optical imaging) are highly complementary to each other, and can be combined together in future studies. Second, in the current study we applied six conditions to demonstrate the effect of CBV on glymphatic system. Given the wide range of pharmacological effects of those anesthetics and caffeine, even more conditions will be helpful. Despite these limitations, the current study provides a unique perspective on the relationship between CBV and glymphatic function.

In conclusion, we have demonstrated that CBV was negatively correlated to the glymphatic function, through altering the extra-ventricular CSF volume. Therefore, properly regulating cerebral blood volume may have the potential to enhance brain waste clearance, which can be beneficial in a wide range of physiological and pathological conditions.

## Materials and Methods

### 1. Animals

All animal experiments were approved by Animal Care and Use Committee of Institute of Neuroscience, Chinese Academy of Sciences, Shanghai, China. Male adult C57BL/6 mice (aged 8-12 weeks and weight between 25 and 30 g) were group-housed in the standard laboratory condition under a 12-h light/dark cycle (light on from 7 a.m. to 7 p.m.) with food and water ad libitum.

### 2. DCE-MRI experiment

#### 2.1. Surgical procedure for DCE-MRI

Anesthetized mice were fixed in a stereotaxic apparatus. Then the scalp was removed, and the exposed skull was cleaned. The tissue adhesive (3M Vetbond™) was applied to the incision edge of skin for protecting wound surface. Afterwards, the head was tilted to form an angle of 120° to the body. Using the occipital crest as a reference point, an almond shaped piece of skin (approximately 1cm) was cut along the midline. The neck muscles were bluntly separated and retracted laterally using 7-0 nylon suture. Then the cisterna magna was visible and a customized borosilicate capillary (tip diameter of approximately 100-150 μm) attached to PE-10 tubing filled with artificial CSF (aCSF) was inserted into the cisterna magna. The cyanoacrylate glue was applied onto the dural membrane and cured by the glue accelerator. Then the dental cement was used to cover the wound and the additional cyanoacrylate glue was dropped to surface of the dental cement for fixing the tubing. The end of the tubing was closed by the mixture of the cyanoacrylate glue and light-curing flowable resin. Afterwards, the head was placed in a horizontal position and the customized DCE-MRI-compatible head holder was implanted (Fig. S1A). Other parts of the exposed skull were covered with a smooth and thin layer of dental cement to avoid inflammation. After the surgery, the carprofen (5 mg/kg, s.c.) was administrated for analgesia.

#### 2.2. Habituation for awake imaging in the awake, dexmedetomidine and caffeine groups

Habituation procedure for awake imaging was adapted from our previous studies (*17*). Briefly, after seven-day recovery from surgery, mice were habituated for MRI scanning for another seven days. Mice were head-fixed on the customized animal bed (Fig. S1B) with the recorded scanning noise of DCE-MRI. The detailed habituation schedule was listed in Fig. 1A. No reward was given during or after the habituation training.

#### 2.3. Anesthetized/awake condition for DCE-MRI

Mice were divided into six groups: awake, dexmedetomidine (DEX), isoflurane (ISO), isoflurane/dexmedetomidine (ISODEX), ketamine/xylazine (KX) and caffeine groups. Each group consisted of 5∼6 mice.

Anesthetized condition included ISO, ISODEX and KX groups. For ISO group, mice were initially anesthetized and intubated with 5% isoflurane, then maintained with mechanical ventilation using 1.3% ISO delivered by 0.4 ml/min mixed oxygen and air (20% : 80%); for ISO/DEX group, mice were initially anesthetized and intubated with 4% ISO combined with DEX (0.1 mg/kg, i.p.), then maintained with mechanical ventilation using 0.5% isoflurane combined with DEX (0.05 mg/kg/h, s.c.); for K/X group, mice were initially anesthetized and intubated with a mixture of ketamine (100 mg/kg, i.p.) and xylazine (20 mg/kg, i.p.), then maintained with an additional 1/4 of the initial dosage every 40 minutes. Anesthetized mice were used for DCE-MRI experiment 24h after surgery. For intubation in all above three groups, a 22G dispenser needle was inserted into the trachea and secured to the customized MRI compatible connector. During mechanical ventilation, mixed oxygen and air (20%: 80%) were supplied through a ventilator (SAR-1000, CWE, Ardmore, USA).

To optimize ventilator settings for maintaining normal physiological conditions, before DCE-MRI experiment, the left femoral artery catheterization and arterial blood gas analysis were performed. The optimal ventilator settings were determined as follows: inspiration/expiration ratio: 40%: 60%, respiratory rate: 80 beats per minute, tidal volume: 0.038 × body weight (in gram) ml (Fig. S2).

Awake imaging condition included awake, DEX and caffeine groups. For the DEX group, mice were sedated with DEX (0.04 mg/kg, i.p.), and maintained with DEX (0.04 mg/kg/h, s.c.); for caffeine group, mice were injected with caffeine (15 mg/kg, i.p.) and an additional 1/3 of the initial dosage was given every 60 minutes.

Mice were positioned in the customized MRI compatible holder system to minimize head movement during scanning. All animals’ cardiac and respiratory signals were monitored by a monitoring system (SA Instruments, NY, USA). For anesthetized or sedated mice, rectal temperature was also monitored and maintained at 37 ± 0.5LJ via a heating pad.

#### 2.4. DCE-MRI protocol

All MRI acquisitions were performed on a Bruker 9.4 Tesla scanner (Bruker Bio Spin, USA). An 86 mm volume coil was used for transmission and a 4-channel cryogenic phased array mouse head coil (Bruker) was used for receiving.

In DCE-MRI experiment, the intracisternal catheter was connected to a PE-10 tube filled with paramagnetic MR contrast agent (Gd-DTPA, 938 Da, Magnevist, Bayer Pharma AG, Leverkusen, Germany) attached to a 100 μl microsyringe (Hamilton-1700) and microinfusion pump (Harvard Apparatus). The imaging protocol mainly consisted of pre-contrast variable flip angle spoiled gradient echo (VFA-SPGR) sequence for T1-mapping and subsequent DCE-MRI using a standard 3D T1-weighted spoiled gradient echo fast low angle shot (3D-T1W-FLASH) sequence. Both sequences are described in detail below.

Pre-contrast T1 mapping was used for quantitative assessment of brain tissue and CSF volume. VFA-SPGR sequence was performed with the following parameters: repetition time (TR) = 15 ms, echo time (TE) = 3.4 ms, average = 1, matrix = 200 × 150 × 150, FOV = 20 × 15 × 15 mm^3^, resolution = 0.1 × 0.1 × 0.1 mm^3^, bandwidth (BW) = 50 kHz, scanning time = 3 min 58 s. A set of six flip angles (2°, 5°, 10°, 15°, 20°, and 30°) were acquired for T1 mapping.

DCE-MRI images were collected with 3D-T1W-FLASH. Imaging parameters were as follows: TR = 15 ms, TE = 3.4 ms, average = 1, flip angle = 15°, matrix = 200 × 150 × 150, FOV = 20 × 15 × 15 mm^3^, resolution = 0.1 × 0.1 × 0.1 mm^3^, BW = 50 kHz, scanning time = 3 min 58 s. The first three images were taken as baseline images, and Gd-DTPA (12.5 mM) infusion was started at the beginning of the fourth image with a constant rate of 0.8 μl/min for a total volume of 10 μl. From the fourth scan, 3D-FLASH images were acquired continuously for at least 160 min, totaling 40 frames for each study. If Gd-DTPA leaked out of the skull cavity, the animal was excluded.

#### 2.5. DCE-MRI Data Analysis

##### 2.5.1. Analysis of Glymphatic Transport from DCE-MRI

The DCE-MRI data pre-processing procedure for each mouse consist of the following steps: First, the acquired DCE-MRI images were converted into the NIFTI file format and the whole brain was extracted manually using ITK-SNAP (http://www.itksnap.org/). Second, head movement was corrected by rigid-body alignment of each scan to the time-averaged image. Third, the above pre-processed images were then nonlinearly transformed to a mouse brain template (https://www.nitrc.org/projects/tpm_mouse) using SPM (http://www.fil.ion.ucl.ac.uk/spm/).

Fourth, spatial smoothing (Gaussian Kernel with isotropic 0.15 mm full width half maximum) was applied to each image. Finally, six brain regions, which included isocortex, thalamus, hypothalamus, hippocampus, striatum and olfactory bulb (Fig. S7A), were defined by the Allen mouse brain atlas (http://atlas.brain-map.org).

After pre-processing, for each post-contrast image, percentage signal change from baseline of each voxel was calculated according to the following formula:

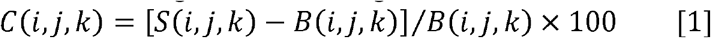

Where C was the signal change percent from baseline, S was post-contrast image signal intensity, B was the averaged signal intensity from three baseline scans, and (i, j, k) was the coordinate of voxel. Then, the averaged signal changes of whole brain and six brain regions at each time point were calculated and time signal curves (TSCs) were obtained.

To evaluate the glymphatic function, we calculated the following parameters: (1) the area under curve (AUC) of TSC was calculated by the conventional trapezoidal integral method (*37*). (2) Arrival time was defined by finding the first time point when signal increased for at least three successive measurements (*38*). (3) The ascending slope was estimated by fitting a line that spans from the baseline to the peak amplitude of signal change from baseline in accordance with previous study (*39*). (4) The number of voxels with enhanced signal was determined at the voxel level. After the administration of Gd-DTPA, if the signal change in a voxel was certain fold (3-fold, 5-fold or 10-fold) higher than specific standard deviations (SD) of baseline signal fluctuations, the signal change in that voxel was considered to be attributed to the arrival of Gd-DTPA.

##### 2.5.2. CSF Segmentation

T1 mapping, exploiting the unique T1 values of gray matter, white matter, and CSF, offers an efficient approach for the segmentation of these distinct tissue types in brain imaging (*40, 41*). When a transverse magnetization is completely spoiled after each RF excitation and the T2* effect is negligibly small (TE≪T2*), the image intensity of VFA-SPGR sequence, S is expressed as

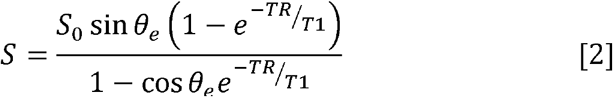

where, θ_e_, TR and S_0_ represent effective flip angle, repetition time and the water proton density weighted signal, respectively. Linearization of Eq. [2] as expressed in Eq. [3] was then solved by an unweighted linear least square fit algorithm to calculate T1 from the slope of the linearized equation (*42*) expressed as

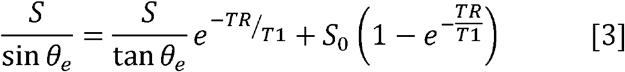

Finally, 3D pre-contrast tissue T1 map was calculated by using the set of six flip angles (2°、5°、 10°、15°、20°、30°) in VFA-SPGR sequence.

T1 Map was then nonlinearly transformed to a mouse brain template for group analysis. Individual T1 maps were segmented into gray matter, white matter, and CSF probability maps using a unified segmentation algorithm in SPM12 using the in vivo mouse tissue probability maps (https://www.nitrc.org/projects/tpm_mouse) as spatially resolved tissue priors. For CSF component, segmented CSF probability maps in standard space were then thresholded at 0.99, 0.9 and 0.8, then the binarized CSF map was further segmented into ventricular and extra-ventricular fractions (Fig. S9) manually by ITK-SNAP. For brain tissue component, segmented gray matter and white matter probability maps were then thresholded at 0, and the binarized gray matter and white matter map collectively represented brain tissue. The volume of each part is obtained by multiplying the number of voxels by the size of the voxels.

### 3. Quantitative cerebral blood volume MRI experiment

#### 3.1 Animal preparation

In this experiment, another group of mice were divided into the same six groups as in the DCE-MRI experiment and had a similar preparation process, except that the cistern magna cannulation was not applied, and contrast agent (Gd-DTPA) was given intravenously. Each group consisted of 6∼9 mice.

#### 3.2. Theory

Quantitative cerebral blood volume is quantified using the improved bookend technique (*21-23*). The concise acquisition and calculation process for quantitative CBV is illustrated in Fig. S5, adapted from the flowchart in the previous study (*23*). This method first calculates relative CBV with standard dynamic susceptibility contrast-enhanced MRI (DSC-MRI) technique, then calibrates the relative CBV values with an AIF (arterial input function) -independent measurement of steady-state CBV. The theoretical background is briefly summarized below.

##### Part one: AIF-dependent Measurement of Relative CBV

The relative CBV (rCBV) based on DSC-MRI technique is obtained by Eq. [4] and [5]:

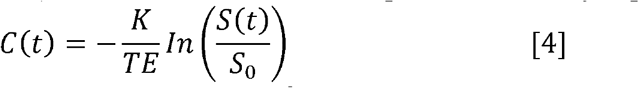

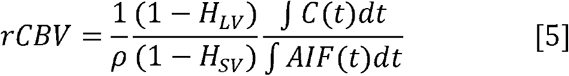

Where C(t) is the contrast agent concentration in tissue at time t after injection of Gd-DTPA, S(t) is the corresponding signal intensity at time t, S_0_ is the pre-injection signal, TE is the echo time, and K reflects the contrast agent relaxivity and properties of the pulse sequence. AIF(t) is the tissue-concentration curve measured in or near an artery, ρ is the density of brain tissue (1.04 g/mL), and H_LV_ and H_SV_ are the hematocrits of the large and small vessels (*21*), respectively.

Considering errors in the measurement of AIF increase the inaccuracy in CBV measurements in DSC-MRI, a calibration based on an AIF-independent steady-state CBV was applied for decreasing the errors introduced by the AIF.

##### Part two: AIF-Independent Steady-state Measurement of CBV

The steady-state CBV (CBV_SS_) is calculated from T1 changes in the white matter (WM) and the large blood vessel, such as superior sagittal sinus (SSS), measured before and after the administration of contrast agent, under the fast exchange approximation:

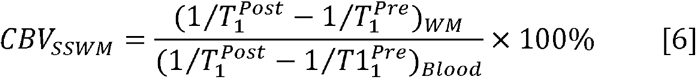

Where 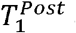 and 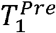 are T1 values before and after contrast agent injection, respectively. Quantitative CBV (in mL/100g) in the WM is then calculated by Eq. [7]:

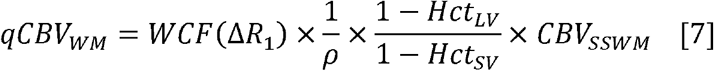

where qCBV_WM_ is the quantitative CBV value in the WM, ρ is the density of brain tissue (1.04 g/mL), H_LV_ and H_SV_ are the hematocrits of the large and small vessels (*21*), respectively, and WCF (ΔR_1_) is the water correction factor determined from the change in 1/T_1_ of blood after contrast injection (*43*), according to the followed Eq. [8]:

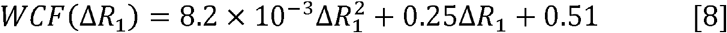

##### Part three: Calculation of quantitative CBV

For the DSC-MRI analysis, the final perfusion quantification in each voxel is calculated with a calibration factor (CF) of WM region using Eq. [9] and [10], where rCBV_WM_ is the rCBV value in the WM.

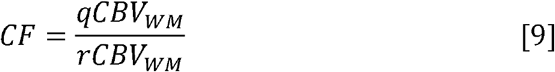

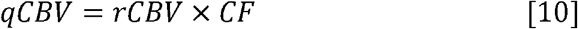

#### 3.3. MRI protocol

To quantify quantitative CBV, a series of MRI sequences were applied in the following order: (1) A 2D inversion recovery spin-echo (IR-SE) sequence was performed to serve as a standard of reference for the inversion recovery fast imaging with steady-state precession (IR-FISP) T1 values. (2) Then, an IR-FISP sequence in a single slice was applied to create a T1 map for calculating CBV_SSWM_ by Eq. [6]. (3) A 2D gradient-recalled single-shot EPI sequence was performed to acquire the perfusion condition during the administration of contrast agent, encompassing 10-s baseline, 6-s injection and 56-s post-injection period. Subsequently, the IR-FISP sequence was repeated. (4) Finally, a 2D time of flight (TOF) FLASH sequence was performed to localize superior sagittal sinus for the CBV_SSWM_ measurement. The detailed sequence parameters as follows:

1. The 2D IR-SE sequence was performed with the following parameters: TR = 6000 ms, TE = 5 ms, BW = 82 kHz, FOV = 16 × 16 mm^2^, matrix size = 100 × 100, slice thickness = 1 mm, slices = 1 (∼ -2.5 mm from bregma with white matter high contrast). Images were acquired at inversion times of 11, 25, 50, 150, 400, 800, 2000, and 5000 ms. The imaging time for each TI was 7 min 36 s, resulting a total of 1 h 44 s.
2. The IR-FISP images were obtained with the following parameter: TR = 2.5 ms, TE = 1.25 ms, FA = 30°, BW = 150 kHz, Inversion Delay = 80 ms, Scan Repetition Time = 8000 ms, segments = 4, Movie Frames = 125, scanning time = 32 s. The FOV, matrix, and prescribed scan location were matched to the IR SE scan.
3. For the perfusion study, the 2D gradient-recalled single-shot EPI images were acquired in the coronal plane with the following parameters: TR = 600 ms, TE = 15 ms, FA = 42°, Dummy Scans = 10, FOV = 16 × 16 mm^2^, matrix = 64 × 64, slice thickness = 0.4 mm, slices = 30, Repetitions = 120. The contrast agent (Gd-DTPA, 0.1 mmol/kg, i.v.) was injected at 5 μl/s, 10 seconds after the scan was initiated. Upon completion of the contrast-enhanced perfusion scan, the IR-FISP scan was repeated.
4. The 2D TOF FLASH sequence was applied with the following parameters: FOV = 16 × 16 mm^2^, matrix = 100 × 100, slice thickness = 1 mm, slices = 1, TR = 15 ms, TE = 3.8 ms, FA = 80°, Averages = 4. The scan location of TOF was identical with the IR-FISP sequence.

#### 3.4. Calculation of quantitative CBV

##### 3.4.1 Determination of the AIF for DSC-MRI

An automated AIF algorithm was applied to identify the voxels that optimally represented the AIF (*44*). The algorithm could be delineated with the following steps. (1) The pre-contrast frames that were used for baseline signal were determined with the whole brain signal intensity as a function of time. The averaged whole brain signal and the SD (σ) were calculated using the first five frames before intravenous contrast injection. The SD was used to determine an upper limit for signal fluctuation to determine pre-contrast frames (< 3σ) and a threshold for contrast agent arrival time of whole brain (> 10σ). (2) By using the pre-contrast frames determined from the analysis of the whole brain, the averaged pre-contrast signal intensity (S_0_) and SD (σ) of each voxel were calculated and used to determine contrast agent arrival time of each voxel (> 5σ). (3) If the arrival time of a voxel was 2 s later than the arrival time of the whole brain, it was assumed to reflect flow in venous structures, which was removed from further AIF analysis. (4) Those voxels (excluding venous voxels) that exhibited the greatest signal depletion and sustained over 4 TRs were defined as AIF voxel. Based on the above AIF voxel, the relative CBV (rCBV) was calculated using Eq. [4] and [5], and rCBV_WM_ was calculated by manually segmenting white matter by ITK-SNAP.

##### 3.4.2. T1 mapping and calibration of IR-FISP T1 values

In this section, CBV_SSWM_ was calculated based on T1 change before and after gadolinium injection. First, the T1 mapping for both IR-FISP and IR-SE sequences was performed using a Reduced-Dimension Non-Linear Least Squares fit algorithm (*45*), as follows:

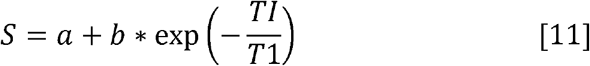

where S was the image signal, TI was the time of inversion, a, b, and T1 were unknown parameters, which needed to be fitted.

Due to the T1 value estimated from IR-FISP sequence was not accurate compared with the T1 value estimated from IR-SE (gold standard), we further used the IR-SE T1 values to calibrate the IR-FISP T1 values. A voxel-by-voxel correlation analysis between IR-FISP and the IR-SE T1 values for each mouse was performed to determine the slope and intercept, which were used to create a calibrated IR-FISP T1 map. Then, the voxels representing the WM were chosen manually segmented by ITK-SNAP, and the voxels of the SSS were defined by a 2D TOF FLASH image. Finally, the CBV_SSWM_ is calculated from the calibrated IR-FISP T1 changes in the WM and the blood vessel measured before and after the administration of contrast agent by Eq. [6]. Next, the quantitative CBV (in mL/100g) in the WM (qCBV_WM_) was calculated by Eq. [7].

##### 3.4.3. Calculation of quantitative CBV

After the CF was calculated by qCBV_WM_ divided by rCBV_WM_ using Eq. [9], the quantitative CBV was calculated using Eq. [10].

Finally, to register the CBV images to standard space, averaged EPI image was first registered to the structure image, then to the standard mouse template. Using the resulting transformation matrix, the CBV images were registered to standard space. ROI-wise mean CBV value was calculated by the corresponding atlas labels (http://mouse.brain-map.org/static/atlas).

### 4. EEG acquisition and analysis

In this section, different mice were used and divided into the six groups, including awake, DEX, ISO, ISODEX, KX and caffeine groups. Each group consisted of 8∼11 mice. Anesthetized mice were fixed in a stereotaxic apparatus. After the above skull cleansing steps, three 0.1 mm diameter holes were drilled in the skull. One gold wire was inserted epidurally at AP = +1 mm, ML= −1 mm to record EEG, and the other two gold wire were implanted into posterior neck muscles to record EMG. Two bare silver wires were used as reference and ground, and inserted epidurally at ML= 0 mm, AP = −5.5 mm and −6.5 mm, respectively. Light curing flowable dental resin was used for fixation of electrodes and head holder. After the surgery, carprofen (5mg/kg, s.c.) was administrated for analgesia.

EEG signals were collected with Synapse software (sampling rate, 1525.9 Hz) outside the magnet. The acquisition time of each mouse was 10 minutes. EEG data were analyzed using a customized MATLAB script. The Chronux toolbox <u>(h</u>t<u>tp://chronux.org/)</u> was used to calculate relative and absolute power of each band (delta, 1 to 4 Hz; theta, 4 to 8 Hz; alpha, 8 to 13 Hz; and beta, 13 to 20 Hz).

### 5. Statistical analysis

Mediation analysis was performed by STATA 17.0 software to examine whether the effect of CBV on glymphatic system function is mediated by the extra-ventricular CSF volume. Specifically, we first estimated the direct relationships between the independent variable (global quantitative CBV) and dependent variable (the AUCs of TSCs). Then, in a mediation model, the extra-ventricular CSF volume was added as the mediator to examine whether full mediation occurs or not. In this context, full mediation occurs when the relationship between the independent variable and the dependent variable is no longer significant with the inclusion of a mediator variable. The significance level was 0.05. A Bootstrap method was performed to evaluate the significance of mediation effect.

Other data were presented as mean ± SD, unless otherwise noted. The statistical analysis was performed using GraphPad Prism 10.0 (https://www.graphpad.com/) or MATLAB 2020b. For the comparison of AUC, arrival time, peak values of signal change, CBV, CSF volume, etc. under six conditions, the one-way ANOVA with Tukey’s correction was used. Linear least squares regression was used for calculation of correlations between group averages. Significance was ascribed at p < 0.05.

## Acknowledgments

The authors thank Yalin Yu (ION, CAS) and Yijuan Zou (ION, CAS) for help in MRI acquisition, and Yue Qiu (Zhongshan Hospital, Fudan University) for help in EEG recording.

## Author contributions

J.C., Z.L. and F.F. designed and supervised the study; J.L. and X.L. performed the animal surgery; J.L., X.L., B.B., M.P., K.Z., M.J., S.Z. and Y.L. collected the electrophysiological and MRI data; J.L., X.L., C.T. and Z.L. conducted the data analysis. J.L., X.L., Z.L. and F.F. wrote the original draft and revised the draft. All authors discussed and commented on the paper.

## Data and materials availability

The MRI data and codes used in this study are available upon reasonable request from the corresponding authors, J.C., Z.L. or F.F..

## References

1. H. Benveniste, The Brain’s Waste-Removal System. Cerebrum : the Dana forum on brain science 2018, (2018).

2. H. Benveniste et al., The glymphatic system and its role in cerebral homeostasis. Journal of applied physiology (Bethesda, Md. : 1985) 129, 1330–1340 (2020).

3. J. J. Iliff et al., A paravascular pathway facilitates CSF flow through the brain parenchyma and the clearance of interstitial solutes, including amyloid β. Science translational medicine 4, 147ra111 (2012).

4. N. A. Jessen, A. S. Munk, I. Lundgaard, M. Nedergaard, The Glymphatic System: A Beginner’s Guide. Neurochemical research 40, 2583–2599 (2015).

5. L. Xie et al., Sleep drives metabolite clearance from the adult brain. Science (New York, N.Y.) 342, 373–377 (2013).

6. C. Gakuba et al., General Anesthesia Inhibits the Activity of the “Glymphatic System”. Theranostics 8, 710–722 (2018).

7. L. M. Hablitz et al., Increased glymphatic influx is correlated with high EEG delta power and low heart rate in mice under anesthesia. Science advances 5, eaav5447 (2019).

8. H. Mestre et al., Flow of cerebrospinal fluid is driven by arterial pulsations and is reduced in hypertension. Nature communications 9, 4878 (2018).

9. J. J. Iliff et al., Cerebral arterial pulsation drives paravascular CSF-interstitial fluid exchange in the murine brain. The Journal of neuroscience : the official journal of the Society for Neuroscience 33, 18190–18199 (2013).

10. S. J. van Veluw et al., Vasomotion as a Driving Force for Paravascular Clearance in the Awake Mouse Brain. Neuron 105, 549–561.e545 (2020).

11. H. Benveniste et al., Anesthesia with Dexmedetomidine and Low-dose Isoflurane Increases Solute Transport via the Glymphatic Pathway in Rat Brain When Compared with High-dose Isoflurane. Anesthesiology 127, 976–988 (2017).

12. H. Mestre et al., Cerebrospinal fluid influx drives acute ischemic tissue swelling. Science (New York, N.Y.) 367, (2020).

13. S. Holstein-Rønsbo et al., Glymphatic influx and clearance are accelerated by neurovascular coupling. Nature neuroscience 26, 1042–1053 (2023).

14. J. R. Goodman, J. J. Iliff, Vasomotor influences on glymphatic-lymphatic coupling and solute trafficking in the central nervous system. Journal of cerebral blood flow and metabolism : official journal of the International Society of Cerebral Blood Flow and Metabolism 40, 1724–1734 (2020).

15. A. S. Thrane, V. Rangroo Thrane, M. Nedergaard, Drowning stars: reassessing the role of astrocytes in brain edema. Trends in neurosciences 37, 620–628 (2014).

16. Z. Han et al., Awake and behaving mouse fMRI during Go/No-Go task. NeuroImage 188, 733–742 (2019).

17. X. Chen et al., Sensory evoked fMRI paradigms in awake mice. NeuroImage 204, 116242 (2020).

18. J. Li et al., Whole-brain mapping of mouse CSF flow via HEAP-METRIC phase-contrast MRI. Magnetic resonance in medicine 87, 2851–2861 (2022).

19. Y. Yu et al., Sleep fMRI with simultaneous electrophysiology at 9.4LJT in male mice. Nature communications 14, 1651 (2023).

20. E. H. Stanton et al., Mapping of CSF transport using high spatiotemporal resolution dynamic contrast-enhanced MRI in mice: Effect of anesthesia. Magnetic resonance in medicine 85, 3326–3342 (2021).

21. K. E. Sakaie et al., Method for improving the accuracy of quantitative cerebral perfusion imaging. Journal of magnetic resonance imaging : JMRI 21, 512–519 (2005).

22. W. Shin, T. A. Cashen, S. W. Horowitz, R. Sawlani, T. J. Carroll, Quantitative CBV measurement from static T1 changes in tissue and correction for intravascular water exchange. Magnetic resonance in medicine 56, 138–145 (2006).

23. W. Shin et al., Quantitative cerebral perfusion using dynamic susceptibility contrast MRI: evaluation of reproducibility and age- and gender-dependence with fully automatic image postprocessing algorithm. Magnetic resonance in medicine 58, 1232–1241 (2007).

24. B. O. Ozturk et al., Disparate volumetric fluid shifts across cerebral tissue compartments with two different anesthetics. Fluids and barriers of the CNS 18, 1 (2021).

25. Y. Chen, T. B. Parrish, Caffeine dose effect on activation-induced BOLD and CBF responses. NeuroImage 46, 577–583 (2009).

26. S. Dreha-Kulaczewski et al., Inspiration is the major regulator of human CSF flow. The Journal of neuroscience : the official journal of the Society for Neuroscience 35, 2485–2491 (2015).

27. S. Dreha-Kulaczewski et al., Identification of the Upward Movement of Human CSF In Vivo and its Relation to the Brain Venous System. The Journal of neuroscience : the official journal of the Society for Neuroscience 37, 2395–2402 (2017).

28. J. K. Holth et al., The sleep-wake cycle regulates brain interstitial fluid tau in mice and CSF tau in humans. Science (New York, N.Y.) 363, 880–884 (2019).

29. K. L. Turner, K. W. Gheres, E. A. Proctor, P. J. Drew, Neurovascular coupling and bilateral connectivity during NREM and REM sleep. eLife 9, (2020).

30. A. Bergel, T. Deffieux, C. Demené, M. Tanter, I. Cohen, Local hippocampal fast gamma rhythms precede brain-wide hyperemic patterns during spontaneous rodent REM sleep. Nature communications 9, 5364 (2018).

31. R. C. Prielipp et al., Dexmedetomidine-induced sedation in volunteers decreases regional and global cerebral blood flow. Anesthesia and analgesia 95, 1052–1059, table of contents (2002).

32. M. M. Todd, J. Weeks, Comparative effects of propofol, pentobarbital, and isoflurane on cerebral blood flow and blood volume. Journal of neurosurgical anesthesiology 8, 296–303 (1996).

33. M. H. Wilson, Monro-Kellie 2.0: The dynamic vascular and venous pathophysiological components of intracranial pressure. Journal of cerebral blood flow and metabolism : official journal of the International Society of Cerebral Blood Flow and Metabolism 36, 1338–1350 (2016).

34. L. M. Miyakoshi et al., The state of brain activity modulates cerebrospinal fluid transport. Progress in neurobiology 229, 102512 (2023).

35. J. Shapey, A. Toma, S. R. Saeed, Physiology of cerebrospinal fluid circulation. Current opinion in otolaryngology & head and neck surgery 27, 326–333 (2019).

36. B. Demiral Ş et al., Apparent diffusion coefficient changes in human brain during sleep - Does it inform on the existence of a glymphatic system? NeuroImage 185, 263–273 (2019).

37. J. J. Iliff et al., Brain-wide pathway for waste clearance captured by contrast-enhanced MRI. The Journal of clinical investigation 123, 1299–1309 (2013).

38. E. Davoodi-Bojd et al., Modeling glymphatic system of the brain using MRI. NeuroImage 188, 616–627 (2019).

39. L. Li et al., Aging-Related Alterations of Glymphatic Transport in Rat: In vivo Magnetic Resonance Imaging and Kinetic Study. Frontiers in aging neuroscience 14, 841798 (2022).

40. H. Lee et al., Quantitative Gd-DOTA uptake from cerebrospinal fluid into rat brain using 3D VFA-SPGR at 9.4T. Magnetic resonance in medicine 79, 1568–1578 (2018).

41. Y. Xue et al., In vivo T1 mapping for quantifying glymphatic system transport and cervical lymph node drainage. Scientific reports 10, 14592 (2020).

42. E. K. Fram et al., Rapid calculation of T1 using variable flip angle gradient refocused imaging. Magnetic resonance imaging 5, 201–208 (1987).

43. T. J. Carroll et al., Quantification of cerebral perfusion using the “bookend technique”: an evaluation in CNS tumors. Magnetic resonance imaging 26, 1352–1359 (2008).

44. T. J. Carroll, H. A. Rowley, V. M. Haughton, Automatic calculation of the arterial input function for cerebral perfusion imaging with MR imaging. Radiology 227, 593–600 (2003).

45. Y. Shen et al., T1 relaxivities of gadolinium-based magnetic resonance contrast agents in human whole blood at 1.5, 3, and 7 T. Investigative radiology 50, 330–338 (2015).

