## Supplemental Figure for "Cerebral Blood Volume Modulates Glymphatic Influx Through Extra-ventricular Cerebrospinal Fluid Volume"

**This PDF file includes:**

Fig. S1. Training apparatus for imaging awake mice.

Fig. S2. Statistics of arterial blood gas analysis across the awake group and two anesthetized groups with endotracheal intubation.

Fig. S3. Contrast agent distribution at five different time points (16 min, 28 min, 44 min, 60 min and 120 min after CM injection) across six conditions.

Fig. S4. Number of voxels with enhanced signal over time across five conditions.

Fig. S5. Acquisition of quantitative CBV data.

Fig. S6. Cerebral blood volume was negatively correlated to glymphatic influx across five conditions.

Fig. S7. Brain region-wise analysis of tracer distribution across five conditions.

Fig. S8. T1 maps from three views across six conditions.

Fig. S9. Detailed flowchart for probabilistic segmentation of CSF volume.

Fig. S10. CSF probability maps from three views across six conditions.

Fig. S11. Changes in CBV altered whole brain tissue volume and further influenced whole brain CSF volume.

Fig. S12. CSF volume and the relationship between CSF volume and glymphatic influx under two different CSF probability thresholds.

Fig. S13. Statistics of arrival time and peak values of DCE-MRI signals for six brain regions under three conditions.

Fig. S14. The glymphatic influx was independent of consciousness level.

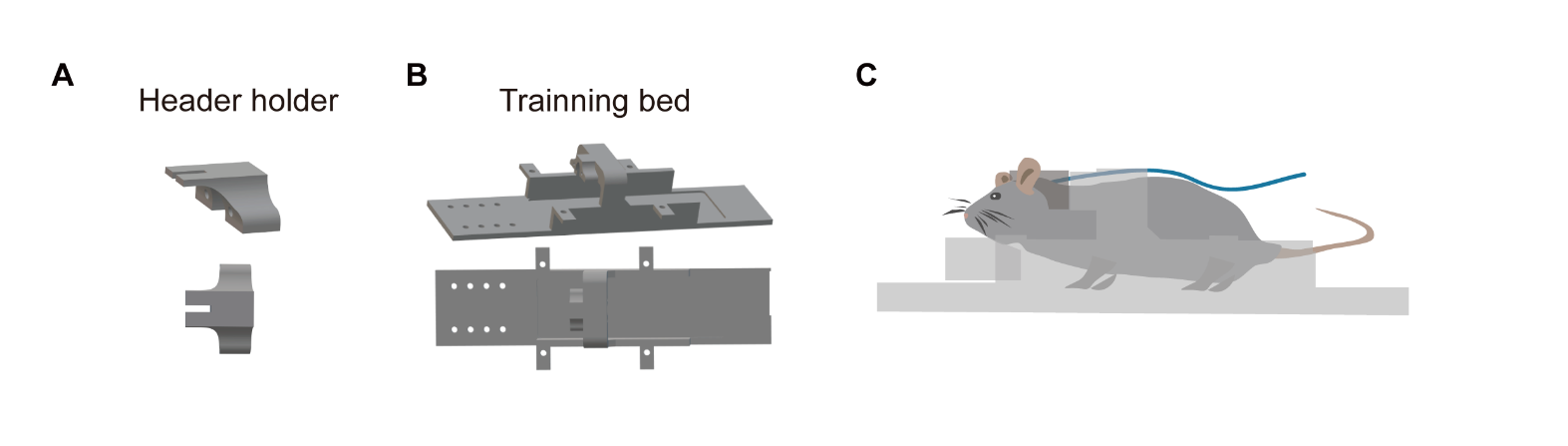

**Fig. S1. Training apparatus for imaging awake mice. (A)** 3D-printed customized head holder used for head fixation. **(B)** 3D-printed customized training bed for mice habituation. **(C)** Schematic diagram of the training process for awake mouse imaging.

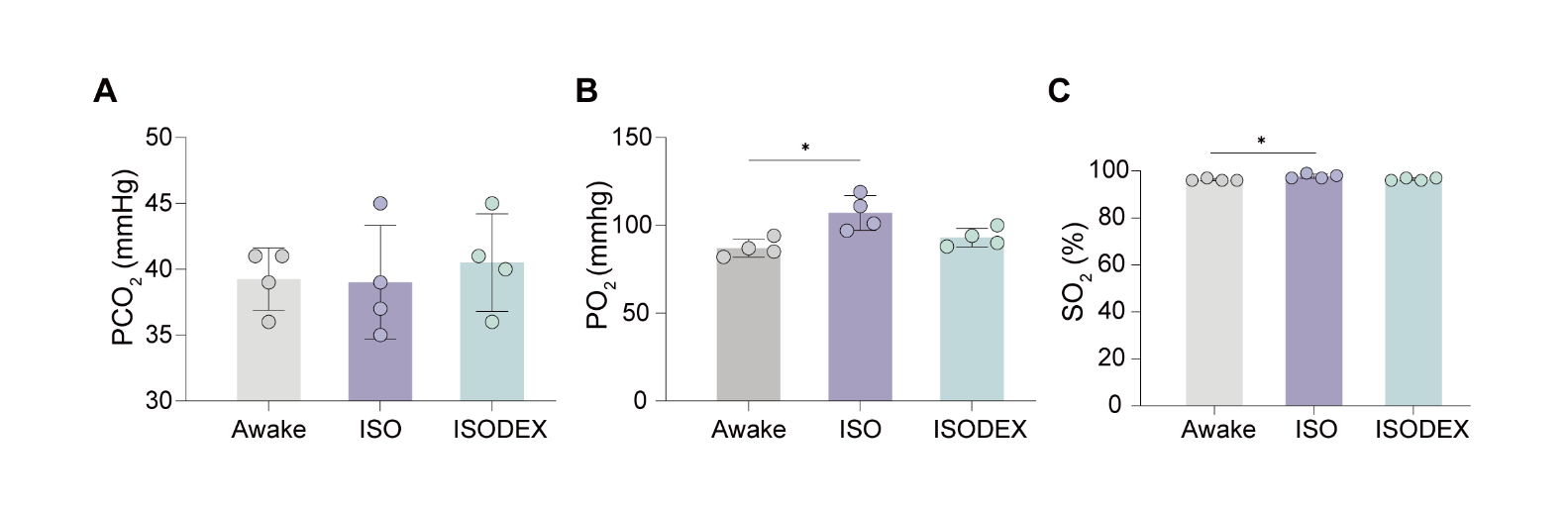

**Fig. S2. Statistics of arterial blood gas analysis across the awake group and two anesthetized groups with endotracheal intubation.** No difference was observed in PCO_2_ **(A)** between awake (n = 4) mice and anesthetized ones under isoflurane (ISO, n = 4) or isoflurane/dexmedetomidine (ISODEX, n = 4) with mechanical ventilation (respiration rate: 80 bpm, tidal volume: 0.038 ml/g). PO_2_ **(B)** and SO_2_ **(C)** levels increased significantly under ISO condition compared with awake condition. These results indicate that blood gas levels are within normal limits in awake and anesthetized endotracheal intubated animals. One-way ANOVA with Tukey’s correction, *, p < 0.05.

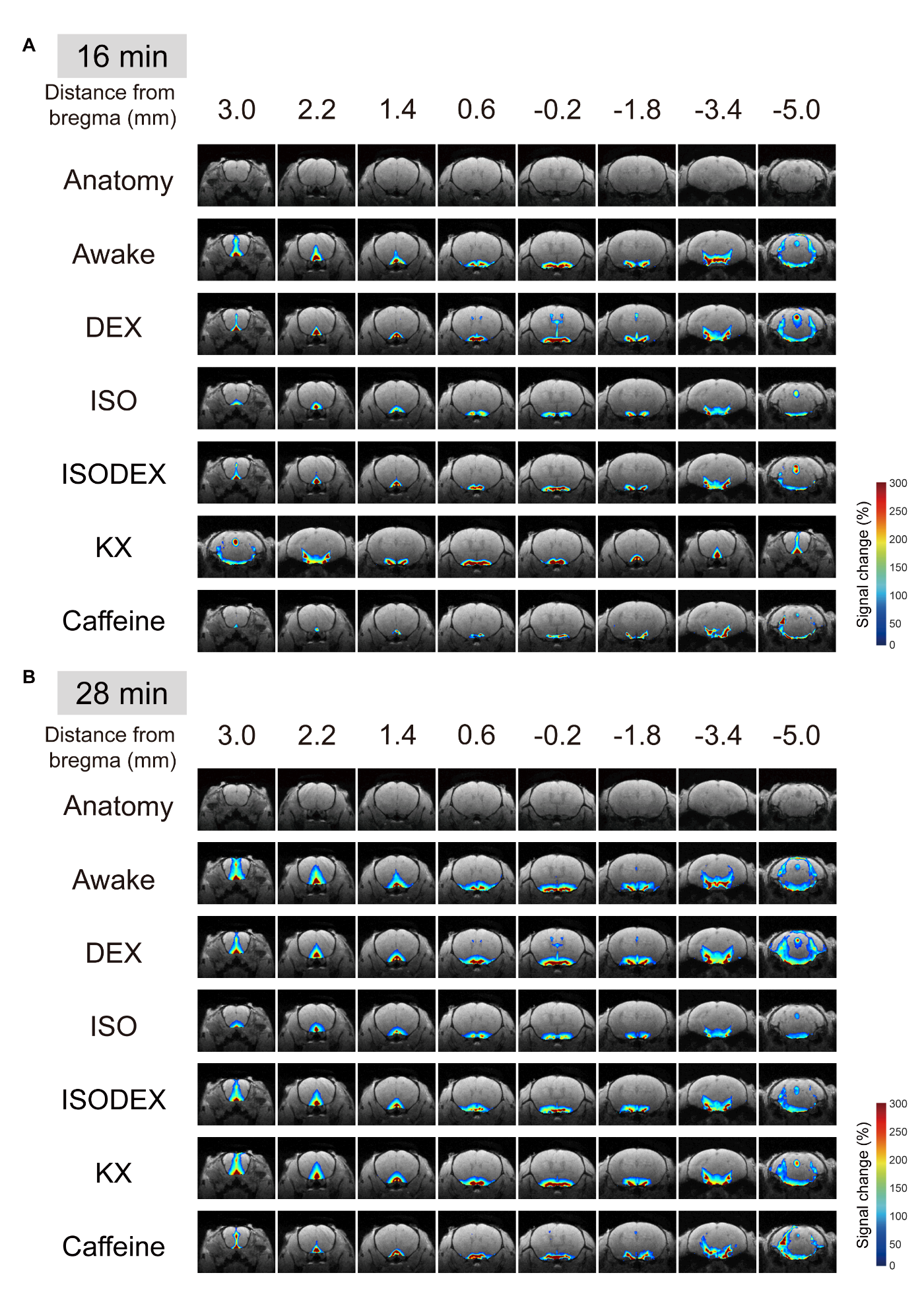

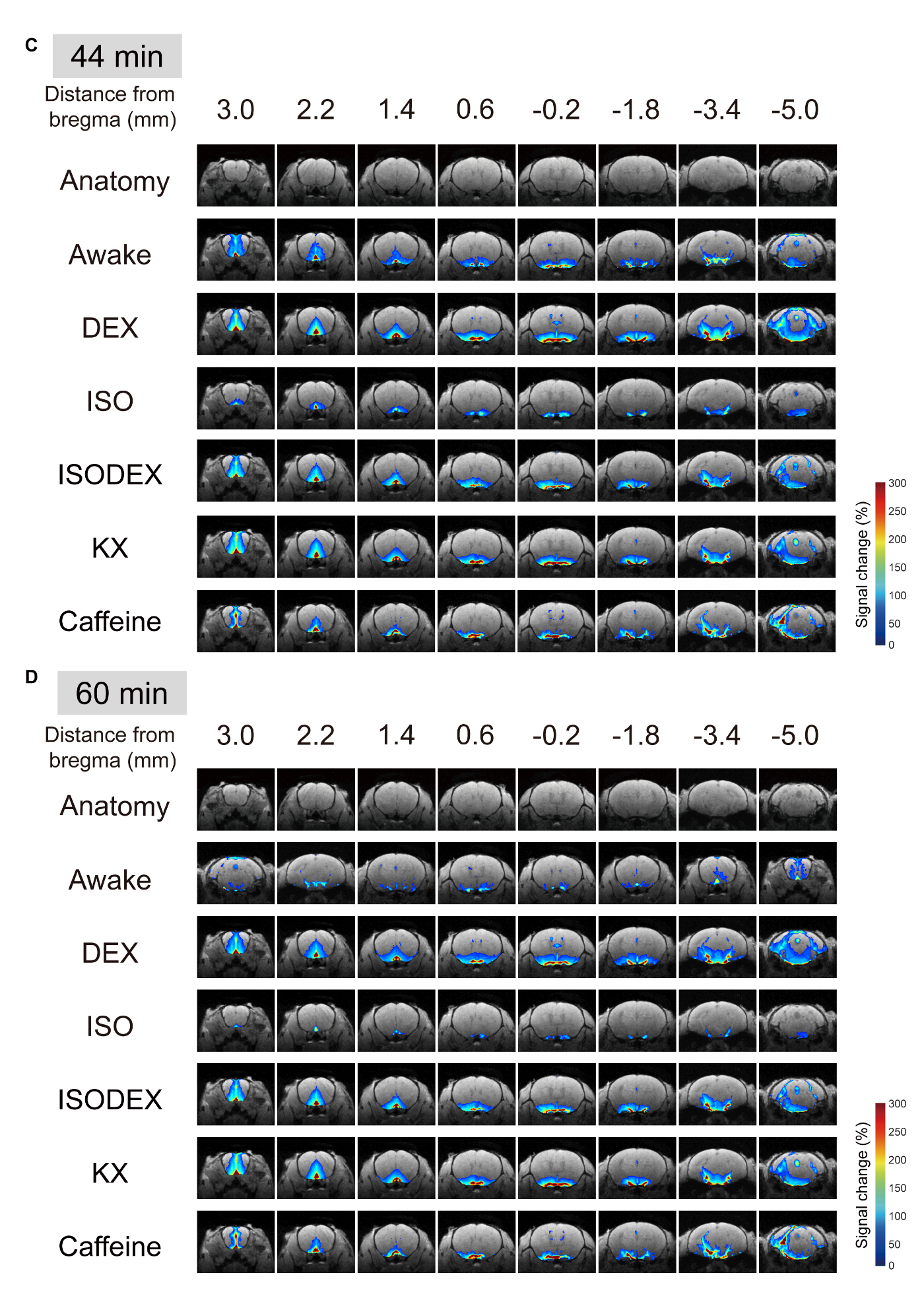

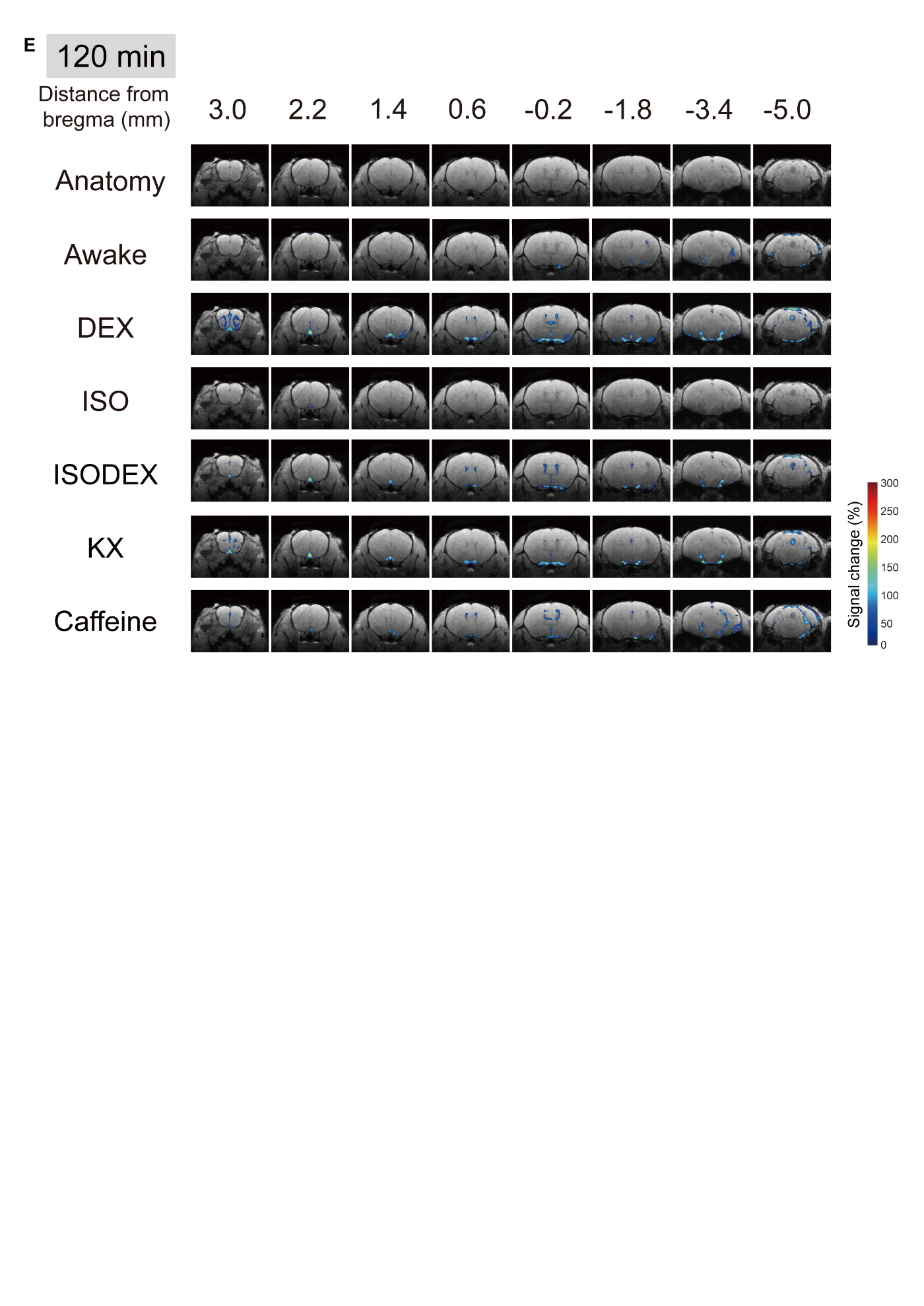

**Fig. S3.** **Contrast agent distribution at five different time points (16 min, 28 min, 44 min, 60 min and 120 min** **after CM injection) across six conditions.** Averaged whole-brain signal change maps in awake (n = 6), DEX (n = 5), ISO (n = 5), ISODEX (n = 5), KX (n = 5) and caffeine (n = 5) groups. **(A–E)** Coronal views of group-averaged pseudocolour-coded DCE-MRI signal change maps, overlaid-with structure images at 16 min **(A)**, 28 min **(B)**, 44 min **(C)**,60 min **(D)** and 120 min **(E)** after injecting Gd-DTPA. The structural images for eight coronal slices at different distances from bregma are labeled in the top row. Six different conditions are labeled in the left column.

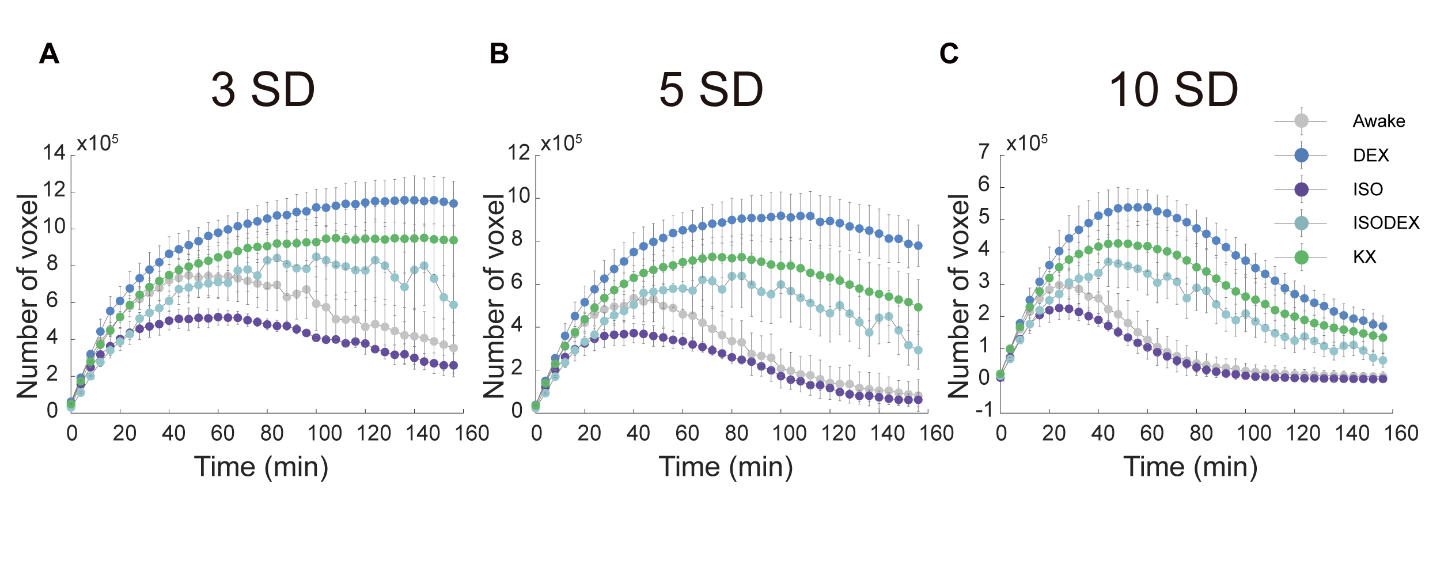

**Fig. S4.** **Number of voxels with enhanced signal over time across five conditions.** Averaged number of voxels with enhanced signal over time in awake (n = 6), DEX (n = 5), ISO (n = 5), ISODEX (n = 5), and KX (n = 5) groups were obtained by setting thresholds at 3-fold **(A)**, 5-fold **(B)** and 10-fold **(C)** above standard deviation (SD) of baseline signal fluctuations. Data are presented as mean ± SEM.

.

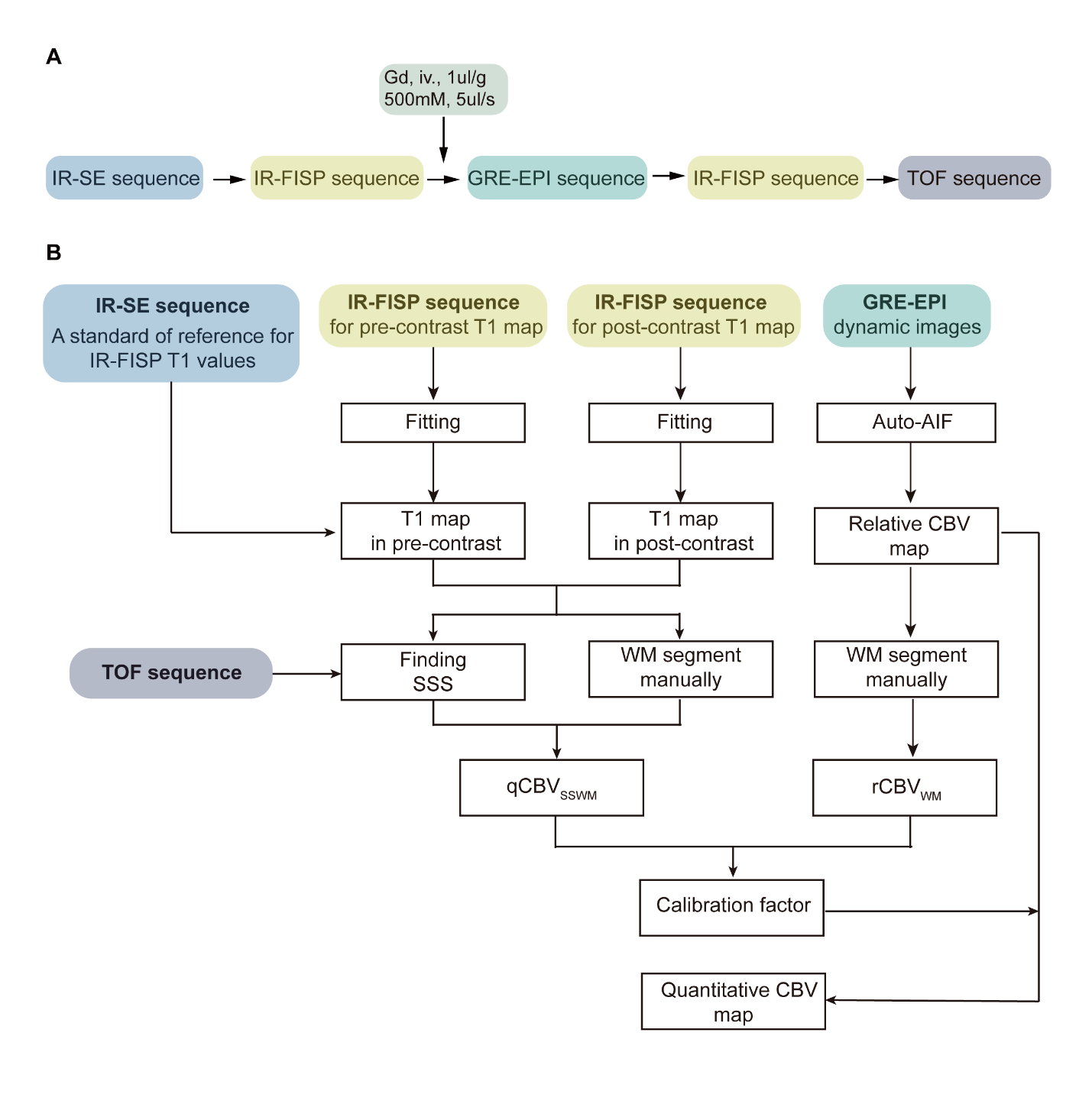

**Fig. S5. Acquisition of quantitative CBV data.** **(A)** Flow diagram of quantitative measurement of CBV using the bookend technique and the sequences included in the process. **(B)** A flowchart of the postprocessing algorithm that calculated quantitative CBV. An averaged CBV value in white matter (qCBV_SSWM_) in steady state was calculated from T1 changes before and after the administration of contrast agent in the white matter (WM) and the large blood vessel, such as superior sagittal sinus (SSS). The calculation of relative CBV (rCBV) was acquired based on an automatically chosen AIF (Auto-AIF). Then, the quantitative CBV was calibrated based on the ratio of qCBV_SSWM_ to relative CBV values in WM (rCBV_WM_).

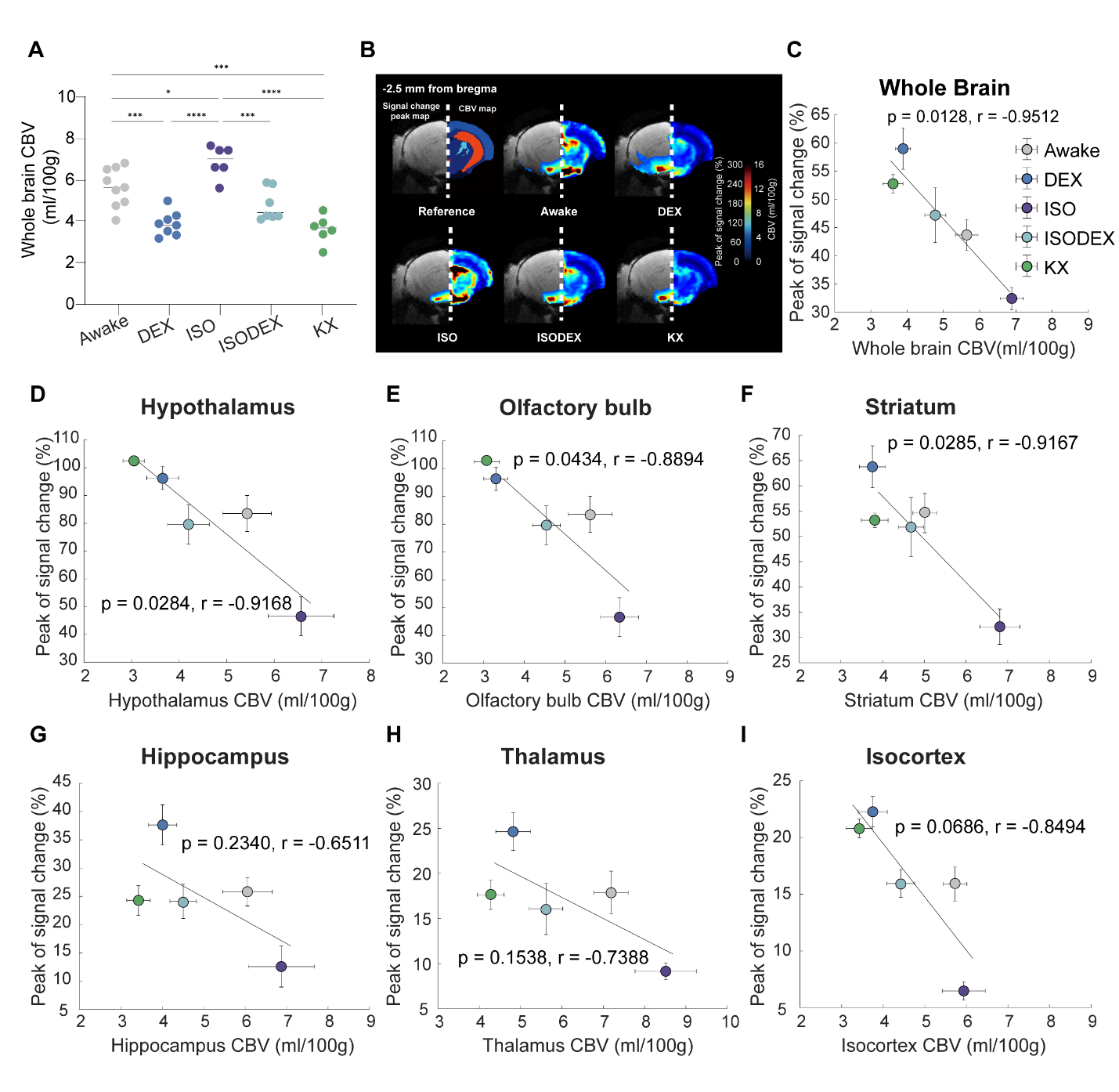

**Fig. S6. Cerebral blood volume was negatively correlated to glymphatic influx across five conditions.** **(A)** The quantitative statistics of whole brain CBV in five groups. One-way ANOVA with Tukey’s correction, *, p < 0.05; ***, p < 0.001; ****, p < 0.0001. **(B)** Population-averaged pseudocolour-coded coronal images (–2.5 mm from bregma) of the Gd-DTPA induced signal change peak maps (left, data from DCE-MRI images) and CBV maps (right). **(C)** Correlation analysis between averaged peak values of TSCs and averaged CBV values from whole brain of five groups. **(D-I)** Correlation analysis between averaged peak values of TSCs and averaged CBV values from hypothalamus **(D)**, olfactory bulb **(E)**, striatum **(F)**, hippocampus **(G)**, thalamus **(H)** and isocortex **(I)** of five groups. Each dot represented the group average (whiskers, SEM).

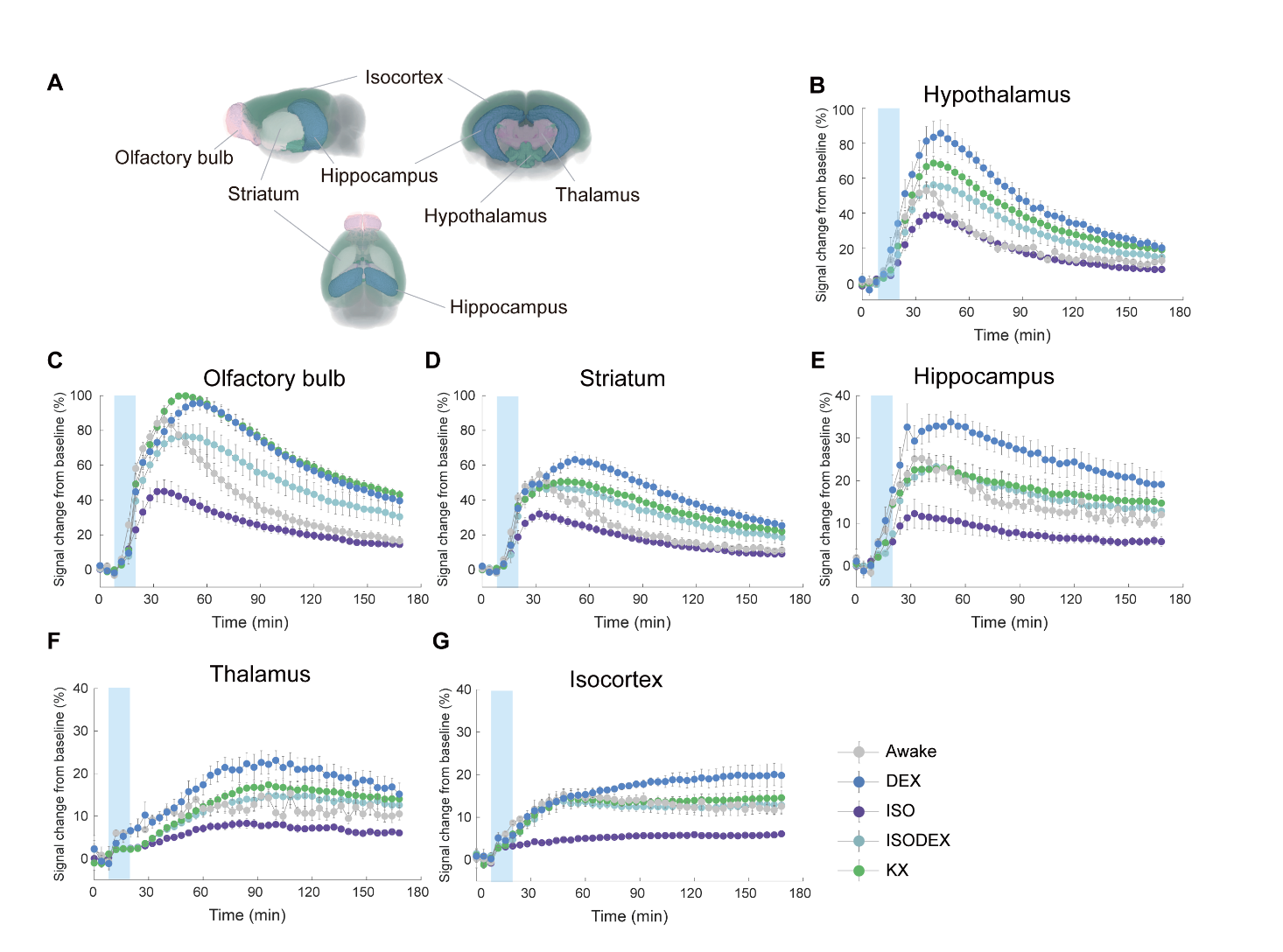

**Fig. S7. Brain region-wise analysis of tracer distribution across five conditions.** Whole brain and region-wise analysis of tracer distribution across five conditions. **(A)** 3D visualization of six regions of interest (ROIs) in the mouse brain including the olfactory bulb, hippocampus, thalamus, hypothalamus, striatum, and isocortex. The averaged TSCs for Gd-DTPA of hypothalamus **(B)**, olfactory bulb **(C)**, striatum **(D)**, hippocampus **(E)**, thalamus **(F)** and isocortex **(G)** were presented. The blue rectangular area represents the contrast agent injection period (12.5 minutes). Data are presented as mean ± SEM.

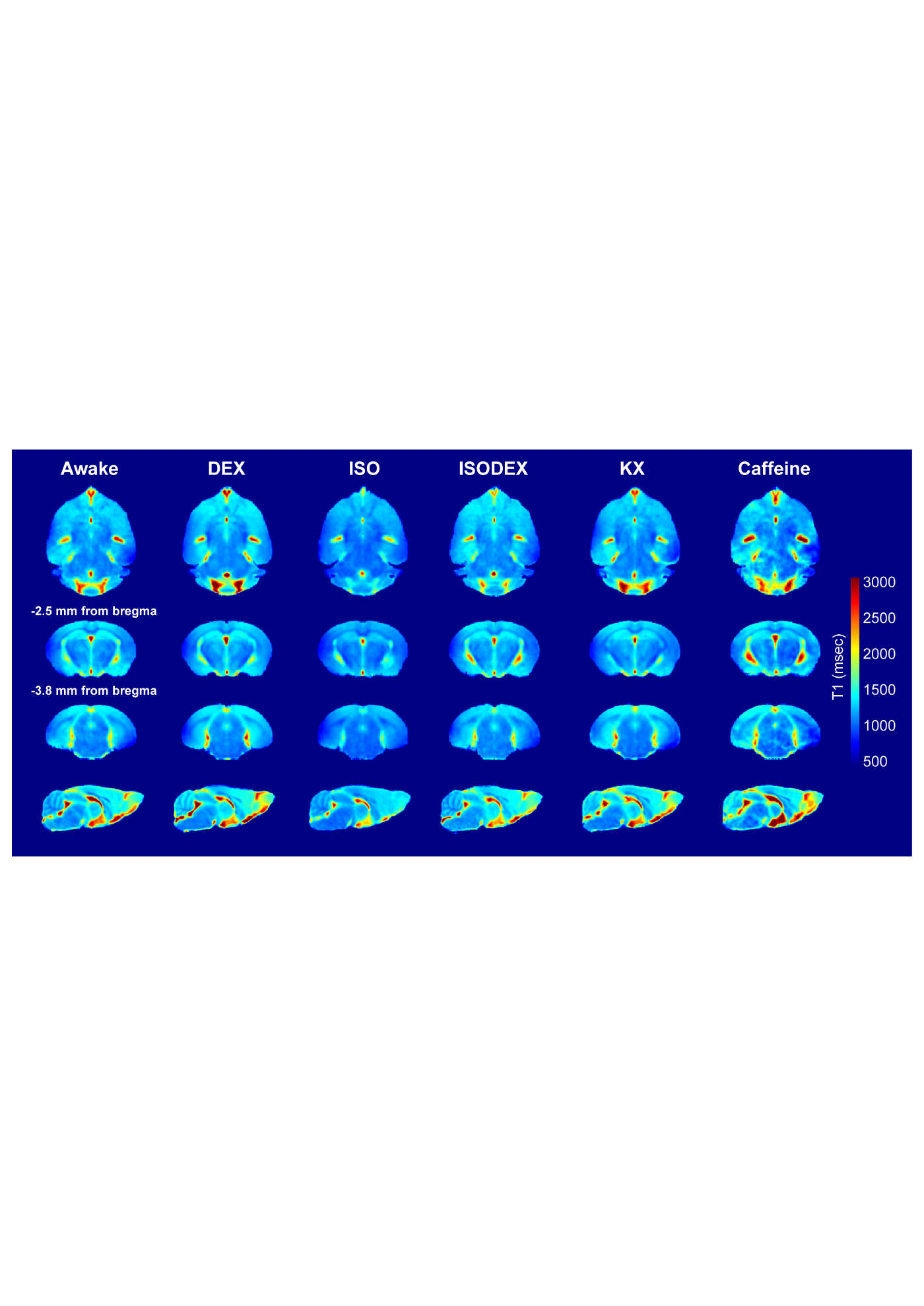

**Fig. S8.** **T1 maps from three views across six conditions.** The pseudocolour-coded averaged T1 maps from the whole brains in three orthogonal planes of mice under awake (n = 6), DEX (n = 6), ISO (n = 7), ISODEX (n = 5), KX (n = 5) and caffeine (n = 5) conditions.

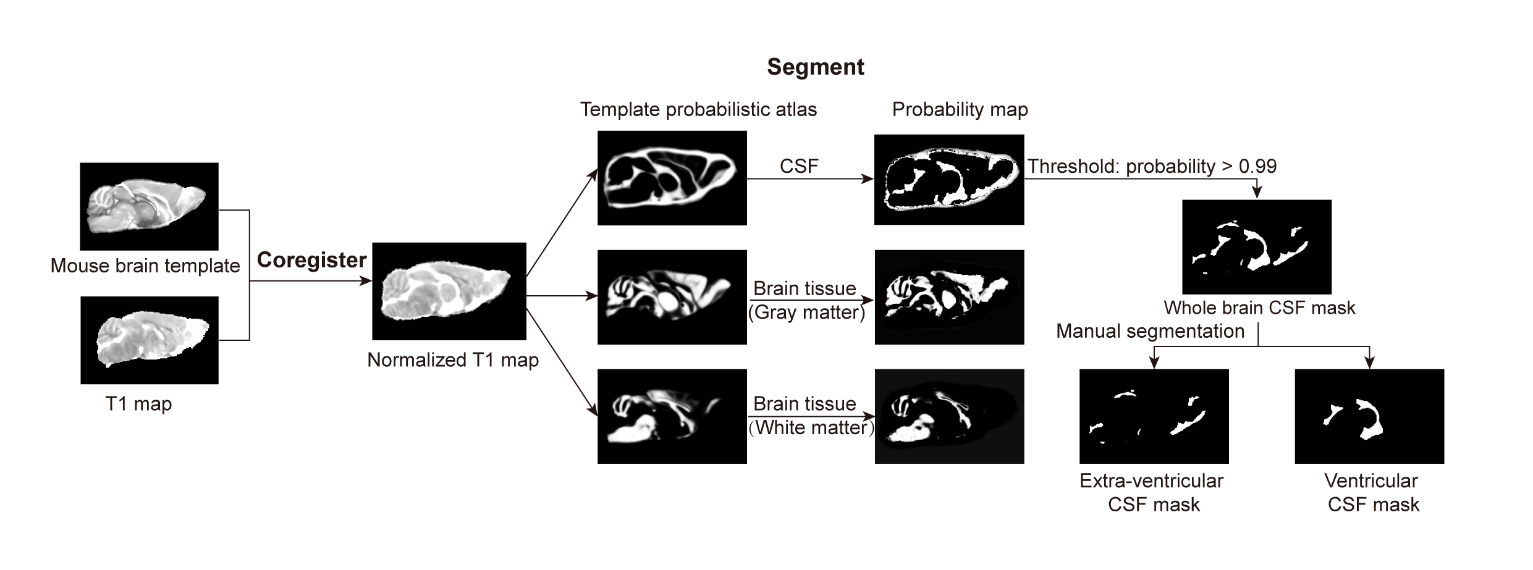

**Fig. S9.** **Detailed flowchart for probabilistic segmentation of CSF volume.** After coregistering the T1 map to the mouse brain template, a tissue probabilistic atlas (including gray matter, white matter, CSF) was used as a spatial prior for normalized T1 map segmentation. After comparing different thresholds (0.8, 0.9 and 0.99) of probability, the CSF mask was created by thresholding the segmented CSF probability map with probability value > 0.99. Finally, manual segmentation was applied to separate the extra-ventricular and ventricular masks.

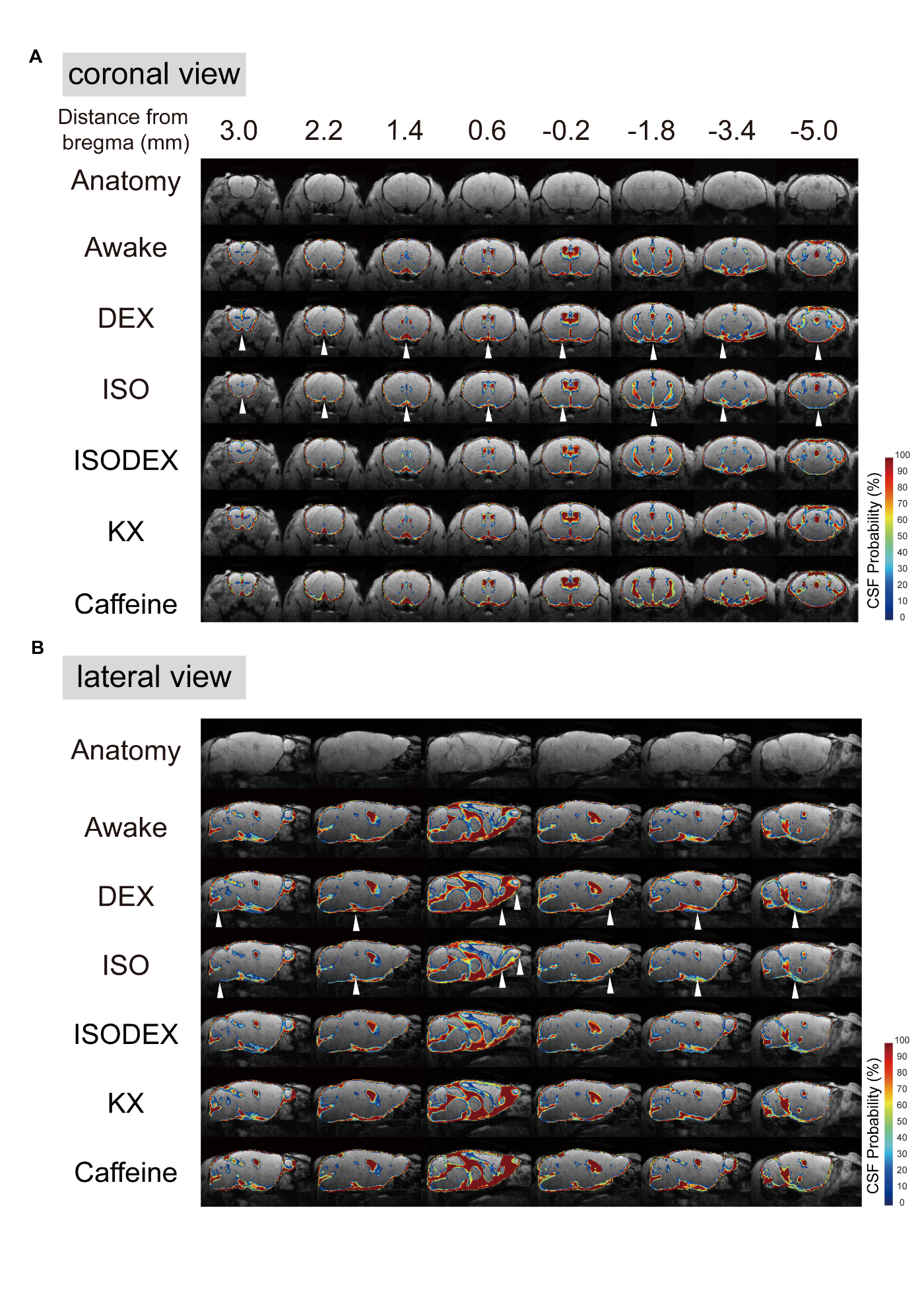

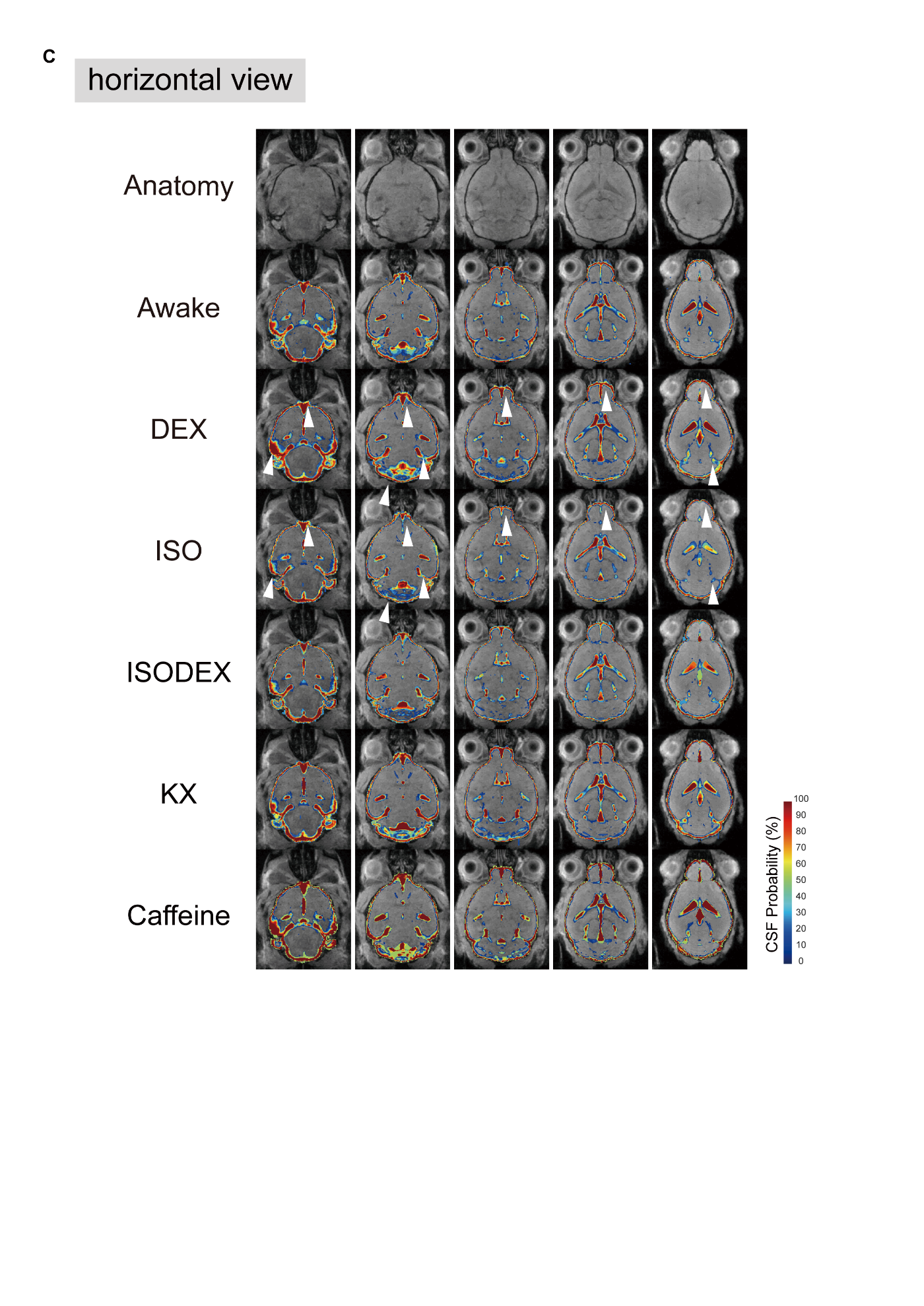

**Fig. S10.** **CSF probability maps from three views across six conditions.** Averaged whole-brain CSF probability maps from three views in awake (n = 6), DEX (n = 6), ISO (n = 7), ISODEX (n = 5), KX (n = 5) and caffeine (n = 5) groups. **(A–C)** Coronal **(A)**, lateral **(B)**, and horizontal **(C)** views of group-averaged pseudocolour-coded CSF probability values, overlaid-with anatomical images. Arrowheads denoted the regions with prominent changes in the CSF probability maps. The anatomical MR from three views images were showed in top row. Numbers, distances from bregma.

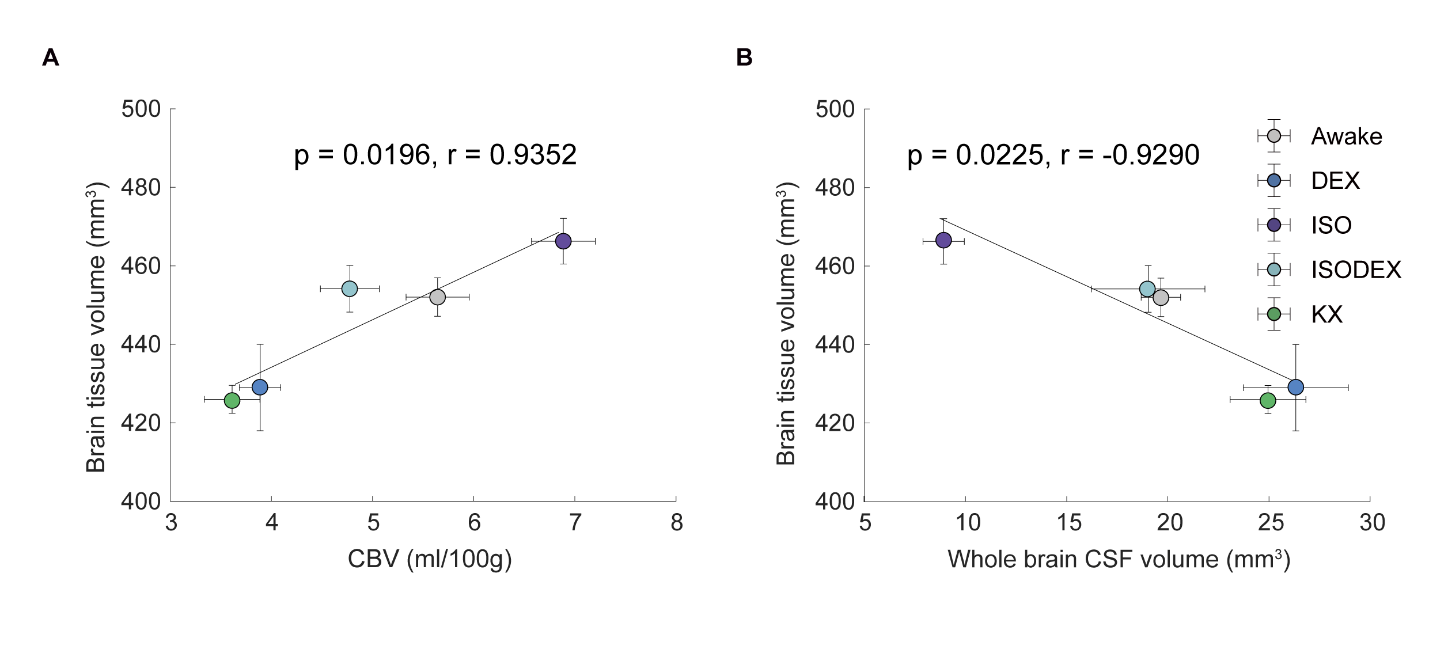

**Fig. S11.** **Changes in CBV altered whole brain tissue volume and further influenced whole brain CSF volume. (A)** Correlation analysis between CBV and brain tissue volume across five conditions. CBV was positively correlated to brain tissue volume across five conditions. **(B)** Correlation analysis between whole brain CSF volume and brain tissue volume across five conditions. Whole brain CSF volume was negatively correlated to brain tissue volume across five conditions. Each dot represented the group average (whiskers, SEM).

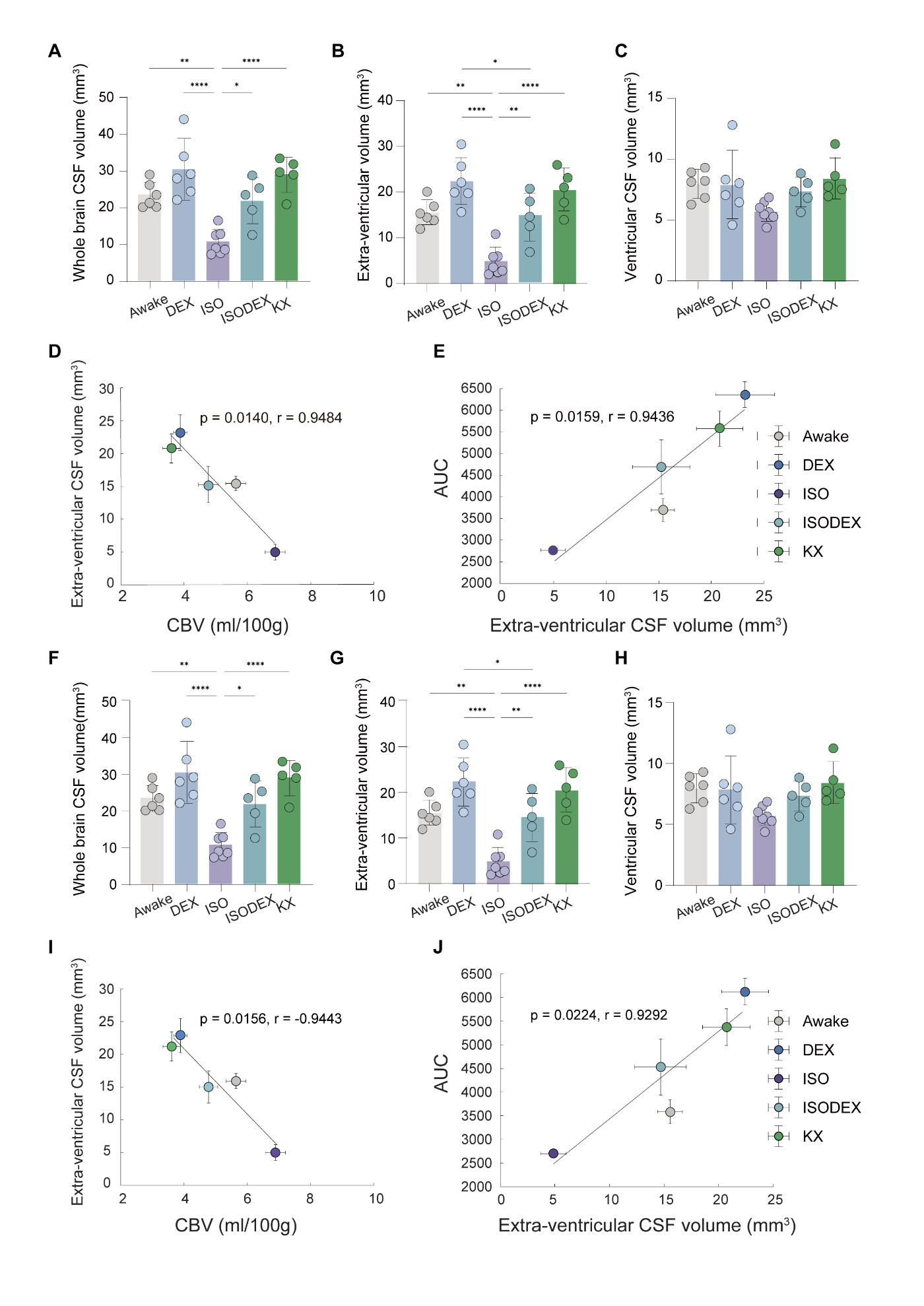

**Fig. S12.** **CSF volume and the relationship between CSF volume and glymphatic influx under two different CSF probability thresholds.** Under the condition that CSF probability threshold was 0.8 **(A-E)** and 0.9 **(F-J)**, statistical comparisons of whole brain CSF volume **(A, F)**, extra-ventricular CSF volume **(B, G)** and ventricular CSF volume **(C, H)** in five groups. One-way ANOVA with Tukey’s correction, *, p < 0.05; **, p < 0.01; ****, p < 0.0001. **(D, I)** Correlation analysis between whole-brain CBV and extra-ventricle CSF volume. Whole-brain CBV was negatively correlated to extra-ventricle CSF volume. **(E, J)** Correlation analysis between extra-ventricular CSF volume and AUCs of whole brain TSCs. Extra-ventricular CSF volume was positively correlated to AUCs of whole brain TSCs. Each dot represented the group average (whiskers, SEM).

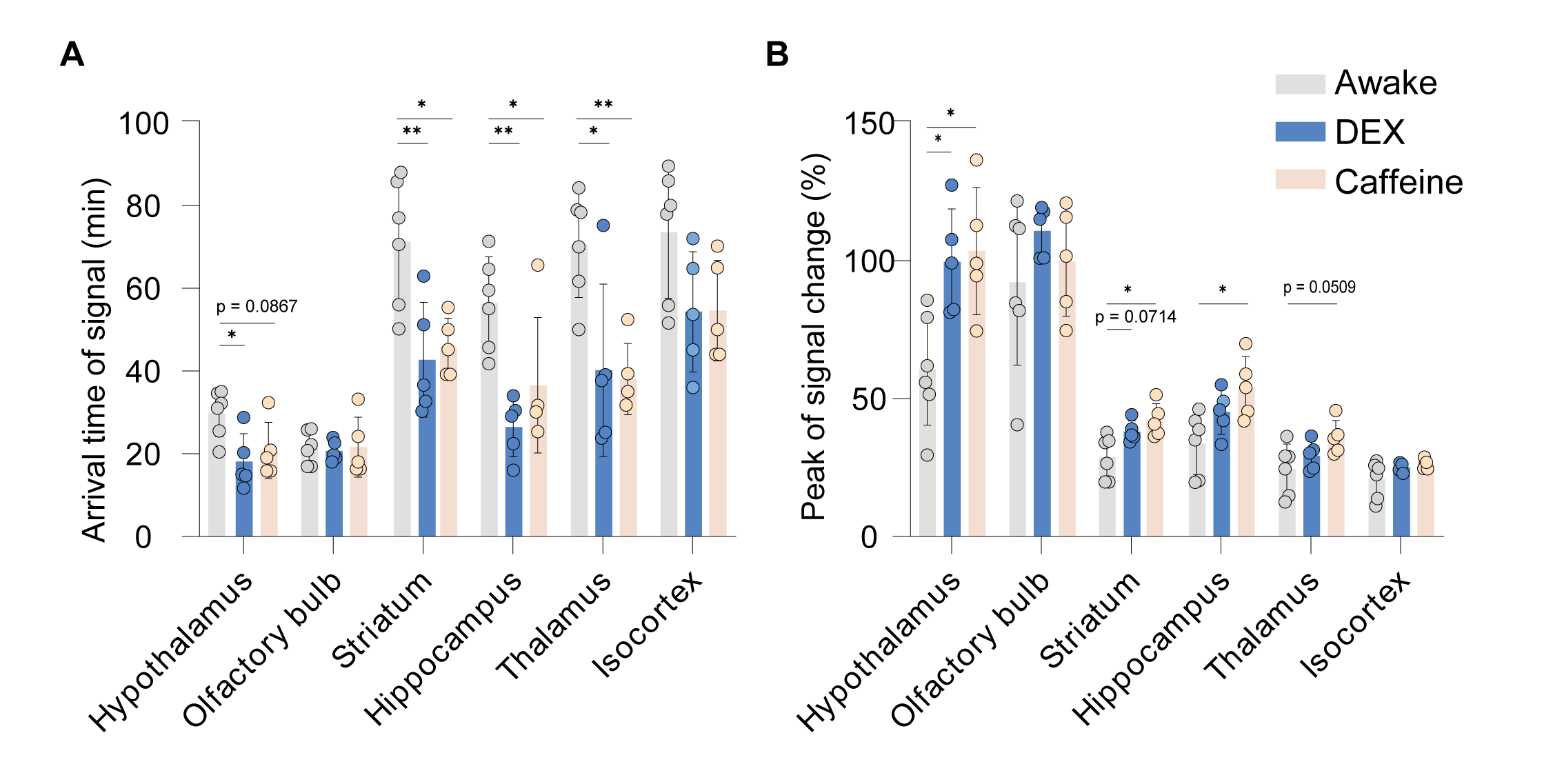

**Fig. S13.** **Statistics of arrival time and peak values of DCE-MRI signals for six brain regions under three conditions.** Statistics of arrival time **(A)** and peak values **(B)** of DCE-MRI signals for hypothalamus, olfactory bulb, striatum, hippocampus, thalamus and isocortex under three conditions. One-way ANOVA with Tukey’s correction, *, p < 0.05; **, p < 0.01.

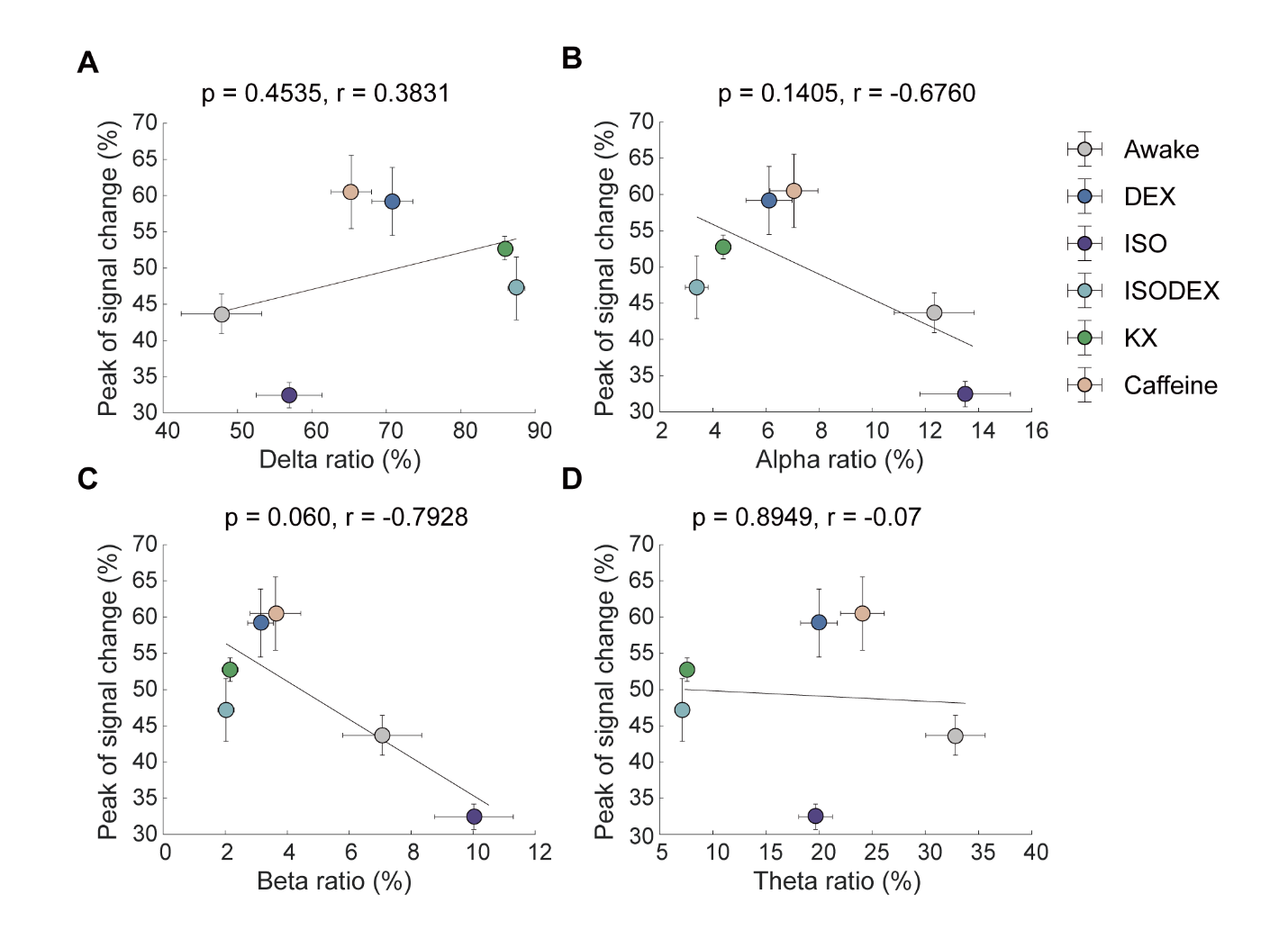

**Fig. S14. The glymphatic influx was independent of consciousness level.** There was no significant correlation between whole-brain peak of signal change and the prevalence of delta **(A)**, alpha **(B)**, beta **(C)**, and theta **(D)** EEG band power across all six groups. Each dot represented the group average (whiskers, SEM). Awake, peak of signal change (n = 6) and EEG (n = 8); DEX, peak of signal change (n = 5) and EEG (n = 10); ISO, peak of signal change (n = 5) and EEG (n = 9); ISODEX, peak of signal change (n = 5) and EEG (n = 10); K/X, peak of signal change (n = 5) and EEG (n = 11); and Caffeine, peak of signal change (n = 5) and EEG (n = 8).
